# High-fidelity Cas13 variants for targeted RNA degradation with minimal collateral effect

**DOI:** 10.1101/2021.12.18.473271

**Authors:** Huawei Tong, Jia Huang, Qingquan Xiao, Bingbing He, Xue Dong, Yuanhua Liu, Xiali Yang, Dingyi Han, Zikang Wang, Wenqin Ying, Runze Zhang, Yu Wei, Xuchen Wang, Chunlong Xu, Yingsi Zhou, Yanfei Li, Minqing Cai, Qifang Wang, Mingxing Xue, Guoling Li, Kailun Fang, Hainan Zhang, Hui Yang

## Abstract

CRISPR-Cas13 systems have recently been employed for targeted RNA degradation in various organisms. However, collateral degradation of bystander RNAs has imposed a major barrier for their *in vivo* applications. We designed a dual-fluorescent reporter system for detecting collateral effects and screening Cas13 variants in mammalian cells. Among over 200 engineered variants, several Cas13 variants (including Cas13d and Cas13X) exhibit efficient on-target activity but markedly reduced collateral activity. Furthermore, transcriptome-wide off-targets and cell growth arrest induced by Cas13 are absent for these variants. Importantly, high-fidelity Cas13 variants show comparable RNA knockdown activity with wild-type Cas13 but no detectable collateral damage in transgenic mice and adeno-associated virus-mediated somatic cell targeting. Thus, high-fidelity Cas13 variants with minimal collateral effect are now available for targeted degradation of RNAs in basic research and therapeutic applications.

## Main Text

The CRISPR and Cas systems have enabled genome editing in various types of cells and organisms^1, 2^. CRISPR-Cas13, the class 2 type VI RNA endonuclease with two Higher Eukaryotes and Prokaryotes Nucleotide-binding (HEPN) domains for RNA cleavage, provides *Escherichia coli* with a programmable immunity against the lytic, single-stranded RNA MS2 bacteriophage^3, 4^, and has been used for RNA manipulations in eukaryotic cells^5–8^. Based on the natural RNase activity on target RNAs, together with collateral cleavage of non-specific single-stranded RNAs *in vitro*, the Cas13 proteins were recently used for nucleic acid detection^9–11^.

The Cas13 protein family was shown to have high efficiency and specificity for programmable RNA targeting, and has been widely used for the cleavage and subsequent degradation of RNAs in yeast, plants, zebrafish, and mammals^5, 7, 12–20^. An ortholog of CRISPR-Cas13d, RfxCas13d (also named CasRx), could mediate RNA knockdown *in vivo* and effectively alleviate disease phenotypes in various mouse models^21–23^. RNA-targeting CRISPR/Cas systems thus provide a promising approach for transcriptome engineering in basic research and therapeutic applications. However, *in vivo* application of CRISPR/Cas13 systems is hindered by the potential existence of collateral effect (Cas13 degrades bystander RNAs once activated by interaction with the target RNA)^3, 24, 25^, one of the fundamental features of CRISPR-Cas immunity^26^. Although collateral RNA degradation by Cas13 has been found in flies^27^, its presence in mammalian cells has not been definitively demonstrated^5–7, 19, 20, 28, 29^. A comprehensive analysis and elimination of collateral RNA degradation are required for *in vivo* applications of Cas13 systems.

Previous structural studies^30–34^ have implicated the mechanisms underlying collateral RNA degradation. Upon binding of Cas13 to target RNA, two HEPN domains undergo distinct conformational changes to form a catalytic site on the protein surface that could degrade both target and non-target RNA at random, a process termed “*cis*/*trans* cleavage” (or “target/collateral cleavage”). However, it is not clear whether there are any distinct binding sites for the target and non-target RNA substrates near the catalytic site of activated Cas13. If so, one may be able to selectively remove the non-target RNA binding through mutagenesis of Cas13, thus eliminating the collateral effect.

We established a rapid and sensitive dual-fluorescence reporter system for detecting collateral effects, and found that Cas13 could induce substantial collateral effects in HEK293T cells when either exogenous or endogenous genes were targeted. Furthermore, we engineered many variants with mutations in HEPN domains, and found several variants with markedly reduced collateral activity and comparable on-target cleavage efficiency. Importantly, we found that the collateral effects revealed by transcriptome-wide RNA-seq and cell proliferation analysis were essentially eliminated when a mutated Cas13 variant Cas13d-N2V8 (967 aa) or Cas13X-M17V6 (775 aa) was used for editing. Thus, by diminishing or eliminating promiscuous RNA binding through mutagenesis, we have obtained several high-fidelity Cas13 variants for targeted RNA degradation with minimal collateral effect.

## Results

### Collateral effects of Cas13 editing in mammalian cells

To evaluate the collateral effect of Cas13 in mammalian cells, we first co-transfected a plasmid coding for EGFP and Cas13a (from *Leptotrichia wadei*, LwaCas13a), or Cas13d (from *Ruminococcus flavefaciens XPD3002*, RfxCas13d), together with a plasmid coding mCherry and guide RNA (gRNA) targeting mCherry (or non-targeting gRNA) into HEK293T cells, and expression levels of EGFP and mCherry were measured 48 hours after transfection (Fig. 1a).

**Fig. 1.**
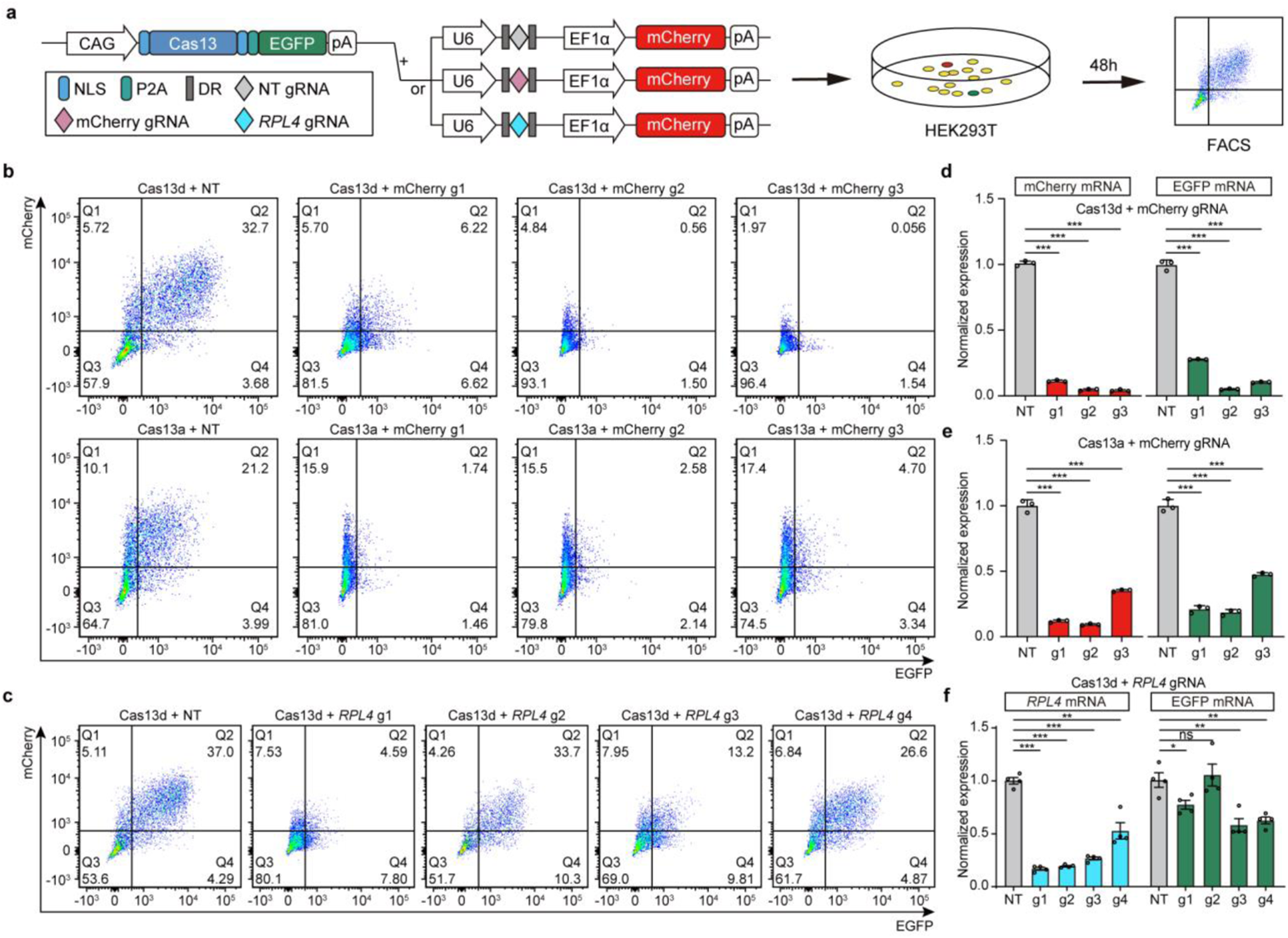
Evaluation of collateral effects in transient transfected HEK293T using dual-fluorescence reporter. **a**, Schematic diagram of the mammalian dual-fluorescence reporter system used to evaluate collateral effects induced by Cas13 (Cas13d or Cas13a)-mediated RNA knockdown. Dual-fluorescence reporter contains one plasmid with Cas13 and EGFP, and one plasmid with gRNA and mCherry. NLS, nuclear localization signal. DR: direct repeat, P2A: 2A peptide from porcine teschovirus-1. **b**, Representative FACS analysis of both mCherry and EGFP fluorescence degradation mediated by Cas13d or Cas13a targeting three different mCherry gRNAs in HEK293T cells, compared with that of non-target gRNA (NT). **c**, Representative FACS analysis of mCherry and EGFP fluorescence degradation induced by Cas13d with four different *RPL4* gRNAs in HEK293T cells, compared with that of non-target gRNA (NT). **d,e**, Relative degradation of mCherry and EGFP transcripts induced by Cas13d (**d**) or Cas13a (**e**) with three different mCherry gRNAs. Degradation relative to a NT gRNA was determined by qPCR, n = 3. **f**, Relative degradation of *RPL4* and EGFP transcripts induced by Cas13d with four different *RPL4* gRNAs. n = 4. All values are presented as mean ± s.e.m.. Two-tailed unpaired two-sample *t*-test. *P < 0.05, **P < 0.01, ***P < 0.001, ns, not significant.

Using fluorescence-activated cell sorting (FACS), we found that co-transfection of mCherry gRNAs (gRNA g1 to g3) with either Cas13a or Cas13d induced significant reduction of not only mCherry fluorescence but also EGFP fluorescence, comparing to the non-targeting (NT) gRNA case (Fig. 1b and Supplementary Fig. 1a-d). The qPCR analysis also showed significant reduction of both EGFP and mCherry transcripts (Fig. 1d,e). These findings revealed substantial collateral effects of Cas13-mediated RNA degradation for targeting transiently overexpressed exogenous genes in HEK293T cells.

Next, the dual fluorescence reporter (EGFP and mCherry) system was used to examine whether Cas13d could induce collateral effects when endogenous genes in HEK293T were targeted. We observed dramatic collateral degradation when targeting the *RPL4* transcript (Fig. 1c and Supplementary Fig. 2a), whereas a slight collateral degradation when targeting *PKM* and *PFN1* transcripts with Cas13d (Supplementary Fig. 2b-f). Furthermore, Cas13d induced robust knockdown of *RPL4* transcript when any one of the four gRNAs (gRNA g1 to g4) was used, and collateral effects as indicated by EGFP reduction were observed for three of the four gRNAs used (Fig. 1f). This observation is consistent with previous reports that the extent of collateral effects differed when different gRNAs were designed for targeting the same gene^28^. This may be attributed to variation of the stability of the activated Cas13/gRNA complexes. Thus, Cas13-mediated RNA degradation results in substantial collateral effects in mammalian cells when targeting either exogenous or endogenous genes.

### Eliminating collateral effects of Cas13 through mutagenesis

To eliminate the collateral effects of Cas13, we sought to engineer Cas13 via mutagenesis and screen for variants free of collateral effects. Firstly, using the dual-fluorescence approach, we constructed an unbiased screening system with EGFP, mCherry, EGFP-targeting gRNA, together with each Cas13 variant into one plasmid, and used FACS to select variants with high-efficiency on-target degradation (based on low EGFP fluorescence) and low collateral degradation activity (based on high mCherry fluorescence) (Fig. 2a). Guided by the structural analysis and biochemical characterization of Cas13^7, 25, 30–34^, we hypothesized that changing RNA-binding cleft proximal to RxxxxH catalytic sites in the HEPN domains may selectively reduce promiscuous RNA binding and collateral degradation while maintaining on-target RNA degradation (Fig. 2b). Thus, we designed and generated a mutagenesis library of over 100 Cas13d variants, each containing mutations in one of 21 segments (N1-N21) (Fig. 2c).

**Fig. 2.**
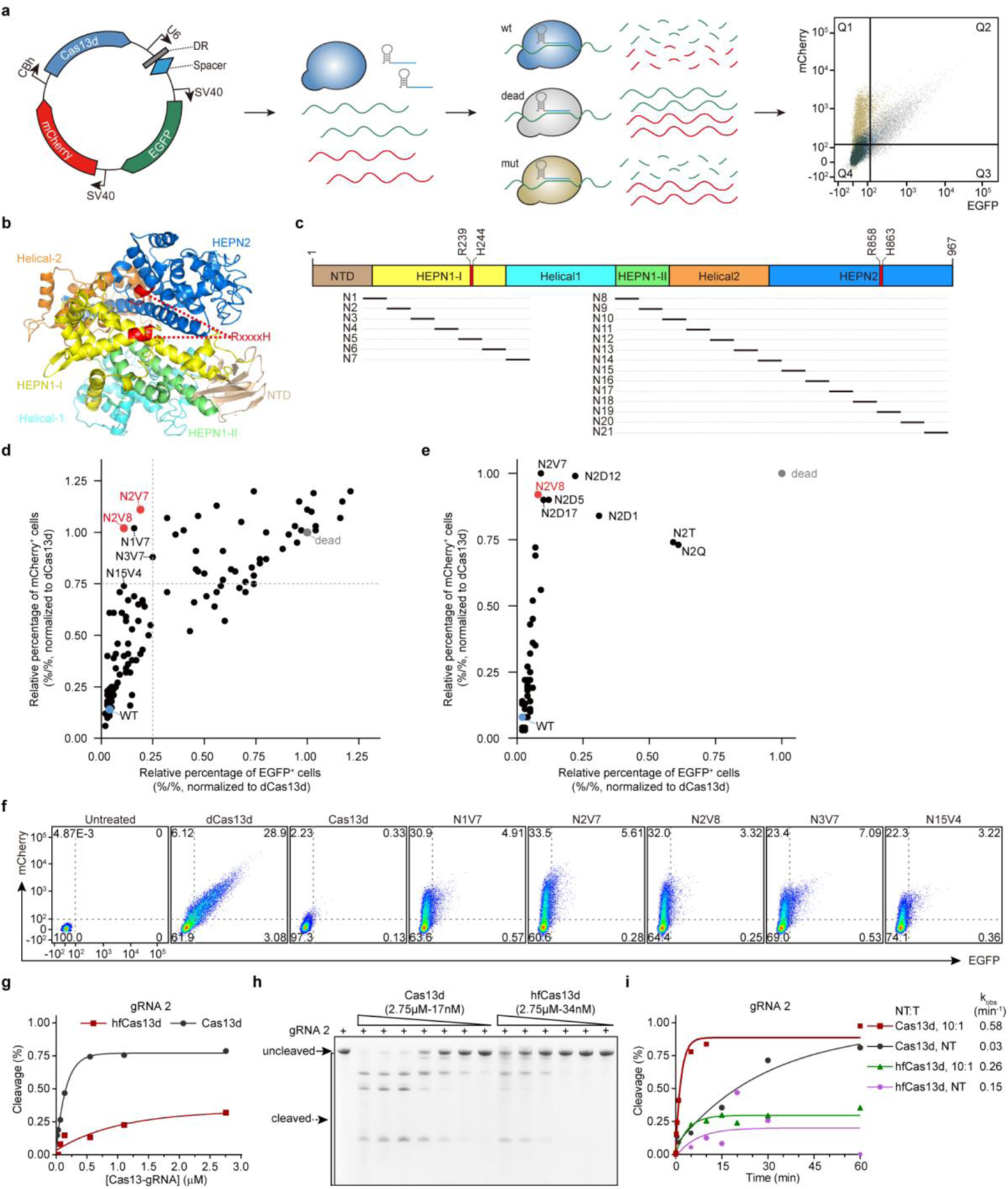
Rational mutagenesis of Cas13d to eliminate collateral activity. **a**, Schematic diagram of the dual-fluorescence reporter system, which contains Cas13d, EGFP, mCherry and EGFP-targeting gRNA in one plasmid. FACS analysis was performed to select mutant variants with low percentage of EGFP^+^ cells and high percentage of mCherry^+^ cells. **b**, View of predicted RfxCas13d structure in ribbon representation. RxxxxHs motifs define the catalytic site, shown as red. **c**, HEPN-1, HEPN-2, Helical-2, and partial Helical-1 domains were selected and divided into 21 segments. **d**,**e**, Quantification of relative percentage of EGFP and/or mCherry positive cells among 118 initially screened Cas13d mutants (**d**) and mutants with different combinations of mutation sites (**e**). WT (wild-type) and dead Cas13d (dCas13d) as controls, relative percentages of fluorescence positive cell were all normalized to dCas13d. Each dot represents the mean of three biological replicates of every mutant variant. **f**, Representative FACS analysis of Cas13d variants with EGFP-targeting gRNA. **g**, Quantification of collateral cleavage activity of Cas13d or hfCas13d with gRNA 2 in the presence of varying concentrations of Cas13:gRNA complex. Exponential fits are shown as solid lines. **h**, Representative denaturing gel depicts cleavage reactions incubated at 37°C for 15 min by Cas13d or hfCas13d with gRNA 2. **i**, Quantified time-course data of collateral cleavage by Cas13d or hfCas13d with gRNA 2. Exponential fits are shown as solid lines, and the calculated pseudo-first-order rate constants (*k_obs_*, mean) of different non-target (NT) to target (T) RNA molar ratios are shown on the right.

We then transfected these variants individually into HEK293 cells and analyzed the reporter fluorescence by FACS. The inactive dead Cas13d (dCas13d, carrying R239A, H244A, R858A, and H863A mutations in HEPN domains) was used as no degradation control (100%). The reduction in the percentage of fluorescence cells, relative to that for dCas13d, indicated the degradation efficiency of the variants. We found that variants with mutation sites in N1, N2, N3, or N15 segments (particularly N1V7, N2V7, N2V8, N3V7, and N15V4) exhibited relatively low percentage of EGFP^+^ cells but high percentage of mCherry^+^ cells, indicating high on-target activity but low collateral activity (Fig. 2d,f). Based on the predicted structures of Cas13d variants (predicted by I-TASSER^35^), we identified that the mutation sites of various effective variants were mainly located in α-helix proximal to catalytic sites of two HEPN domains (RxxxxH-1, RxxxxH-2) (Supplementary Fig. 3). We also examined the effect of reducing the number of mutations in the N2 region from 4 to 3, 2, or 1, and found that the variant with 4 mutations Cas13d-N2V8 had the highest specificity in targeting RNA for degradation (Fig. 2e). Thus, Cas13d-N2V8, termed hereafter as high-fidelity Cas13d (hfCas13d), was used in following experiments.

We next targeted the EGFP transcript with three additional gRNAs (gRNA, g2 to g4), and found essentially no collateral effect by hfCas13d with gRNA g2 or gRNA g4 (Supplementary Fig. 4). We also performed a standard 1:2 serial dilution to dilute Cas13d or hfCas13d plasmids targeting the EGFP transcript with gRNA g2 in HEK293 cells. Cas13d and hfCas13d showed a similar tendency of on-target degradation activity, while hfCas13d showed largely reduced collateral effects compared with Cas13d (Supplementary Fig. 5). However, hfCas13d with gRNA g3 still induced collateral effect to some extent, although lower than that induced by wild-type Cas13d, indicating that gRNA selection is still necessary to achieve free-of-collateral effect with hfCas13d (Supplementary Fig. 4).

To further confirm that hfCas13d has minimal collateral activity, we performed *in vitro* collateral cleavage assay with purified wild-type Cas13d or hfCas13d protein (Supplementary Fig. 6). With different gRNAs (gRNA 1 to gRNA 5), we found obviously reduced collateral cleavage activity for hfCas13d at different concentrations, comparing with Cas13d in all conditions (Fig. 2g-h and Supplementary Fig. 7). Among 5 gRNAs tested, we found very low collateral cleavage with gRNA-2, gRNA-3 or gRNA-4, even with extremely high concentration (2.75 μM, typically 100 nM) of hfCas13d. We also incubated hfCas13d with gRNA-2 at a high concentration (550 nM) for longer time and still found quite low collateral cleavage (Fig. 2i).

Taken together, compared with wild-type Cas13d, hfCas13d showed remarkably reduced collateral cleavage activity while maintained on-target activity.

### Efficacy and specificity of hfCas13d in mammalian cells

Considering the expression level of EGFP transcripts via transient transfection was much higher than that of endogenous transcripts, we next targeted endogenous genes to examine the collateral effects of Cas13d and hfCas13d in HEK293 cells. To evaluate whether the extent of collateral effects induced by Cas13d was correlated with the expression level of endogenous genes, we selected a panel of 23 endogenous genes with diverse roles and differential expression levels in mammalian cells, and designed 1-7 gRNAs for each gene (Fig. 3a). We transfected HEK293 cells with a construct containing Cas13d, EGFP, mCherry, on-target gRNA for each endogenous gene or non-target gRNA (NT), together with another construct coding for blue fluorescent protein (BFP), which was used for normalizing the transfection efficiency. We examined the EGFP and mCherry fluorescence for the collateral effect 48 hours post-transfection. Overall, we found that the higher the expression level of endogenous target gene, the more the collateral degradation induced by Cas13d (Fig. 3b,c). By contrast, we found no detectable collateral degradation induced by hfCas13d with all tested on-target gRNAs (Fig. 3d,e). To compare the RNA degradation activity of Cas13d and hfCas13d, we transfected Cas13d, hfCas13d or dCas13d, together with gRNAs targeting each transcript in HEK293 cells, and performed qPCR assay for each transcript 48 hours post-transfection. We found both Cas13d and hfCas13d exhibited robust degradation of targeted transcripts as further confirmed by qPCR analysis (Fig. 3f-h), although the efficiency of targeted RNA degradation was slightly decreased in hfCas13d when compared with Cas13d (76 ± 2% and 80 ± 1%, respectively), indicating that hfCas13d retained a high efficiency of RNA degradation on many endogenous genes (Fig. 3f-i). In addition, to achieve RNA degradation of targeted transcripts more efficiently by hfCas13d, we screened several gRNAs targeting the same transcript, and found hfCas13d with specific gRNA (e.g., gRNA-1 for *RPL4*, gRNA-7 for *CA2*) exhibited most efficient degradation of target transcripts while induced no detectable collateral degradation (Fig. 3b,d,f,g and Supplementary Fig. 8). By contrast, Cas13d with different gRNAs exhibited different extent of collateral degradation (Fig. 3b). These results indicate that the collateral effect induced by Cas13-mediated RNA degradation correlates with the expression level of endogenous transcript and is essentially eliminated for mutated variant hfCas13d.

**Fig. 3.**
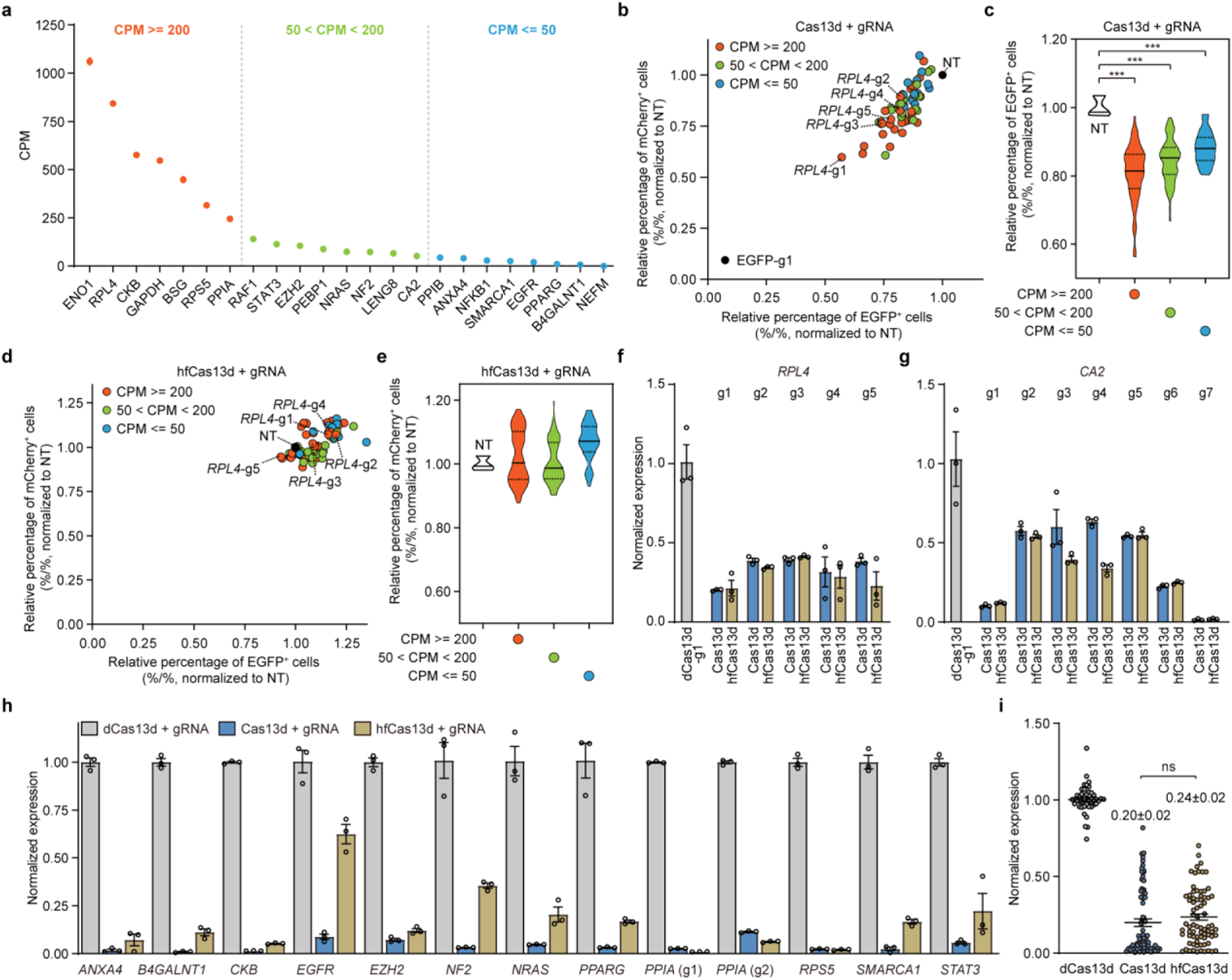
Efficient and specific interference activity of hfCas13d in HEK293 cells. **a**, Relative expression level of 23 endogenous genes in HEK293 cells from RNA-seq of dCas13d groups. Each dot represents mean ± s.e.m. of CPM (Counts per million) for each endogenous gene in dCas13d groups from Figure 4. **b**, Differential reduction of relative percentage of EGFP and/or mCherry positive cells was induced by Cas13d targeting 22 endogenous genes, with 1-7 gRNAs for each transcript, compared with non-targeting gRNA (NT). Each dot represents the mean of three biological replicates for each gRNA. **c**, Statistical quantification from (**b**). **d**, Differential reduction of relative percentage of EGFP and/or mCherry positive cells was induced by hfCas13d. **e**, Statistical quantification from (**d**). **f**,**g**, Relative target RNA knockdown by individual gRNAs targeting *RPL4* (**f**) or *CA2* (**g**). The dCas13d-g1 as vehicle control, respectively. n = 3. **h**, Cas13d and hfCas13d targeting 13 endogenous transcripts. Transcript levels are relative to dCas13d as vehicle control, n = 3. **i**, Statistical data analysis from (**f-h**), each point represents the value of corresponding sample. All values are presented as mean ± s.e.m.. Two-tailed unpaired two-sample *t*-test. *P < 0.05, **P < 0.01, ***P < 0.001, ns, not significant.

### Absence of transcriptome-wide collateral effect for hfCas13d

To comprehensively detect the collateral effect accompanied with targeted RNA degradation by Cas13d and hfCas13d, we performed transcriptome-wide RNA sequencing (RNA-seq) of HEK293 cells 48 hours following transfection with Cas13d, hfCas13d, or dCas13d. We observed widespread off-target transcript degradation in cells expressing Cas13d with *PPIA* gRNA1. Along with high efficiency of *PPIA* on-target degradation, 9289 and 2676 down- and up-regulated genes were significantly changed when compared to dCas13d control, respectively (Fig. 4a,e,f). In addition, we predicted 96 *PPIA* gRNA-dependent off-target genes using blastn-based scripts (see Methods), and found 6 off-target genes down-regulated, most of which are paralogs of *PPIA* (Fig. 4a,g). Furthermore, down-regulation of most of these gRNA-dependent off-target genes could be eliminated when targeting *PPIA* with a different gRNA (Fig. 4b).

**Fig. 4.**
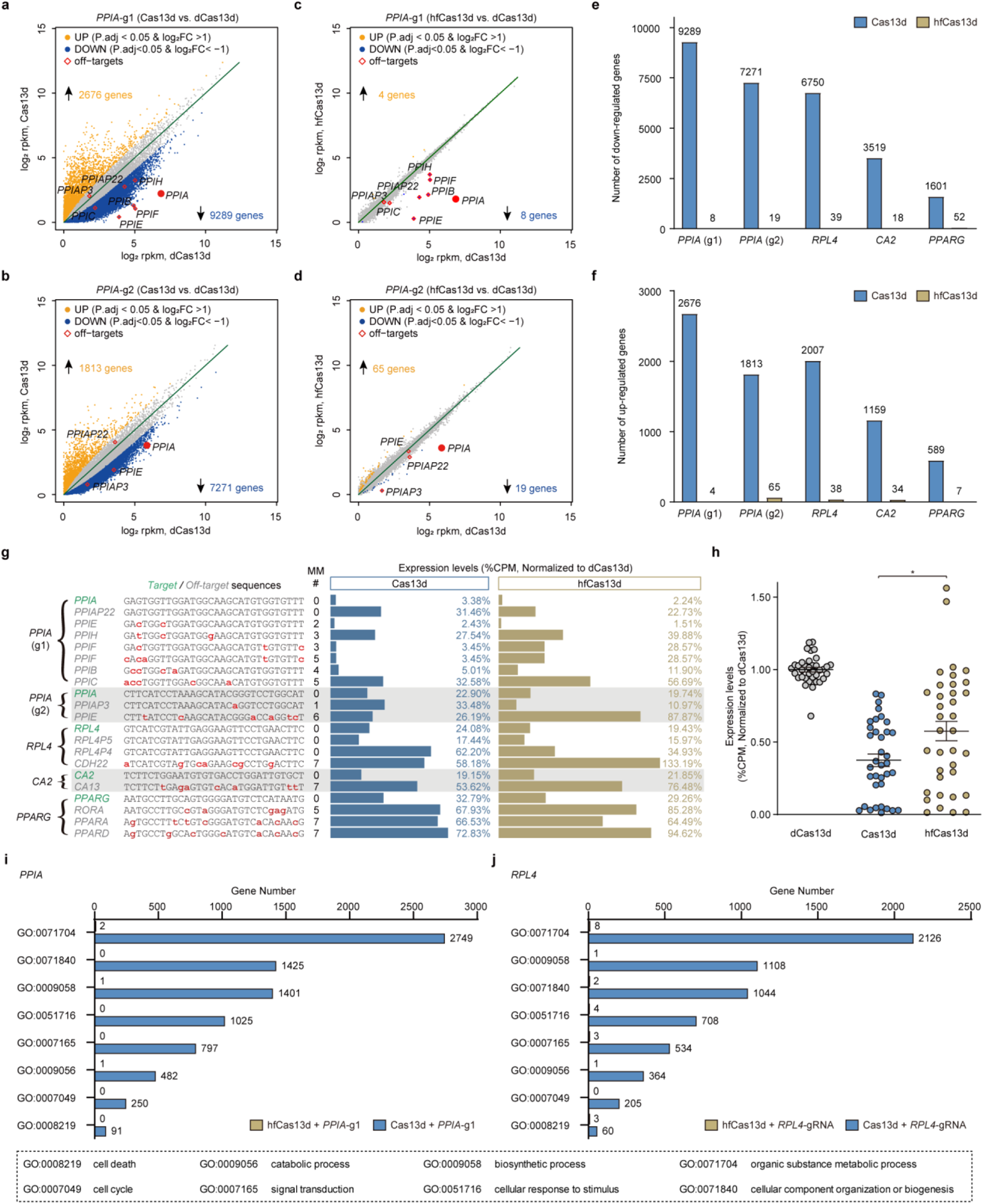
Transcriptome-wide collateral effect analysis for Cas13d and hfCas13d. **a**,**b**, Scatter plot of differential transcript levels between Cas13d- and dCas13d-mediated *PPIA* degradation using gRNA g1 (**a**) or gRNA g2 (**b**). **c**,**d**, Scatter plot of differential transcript levels between hfCas13d- and dCas13d-mediated *PPIA* degradation using gRNA g1 (**c**, n = 2 for hfCas13d; n = 3 for dCas13d) or gRNA g2 (**d**). **e**,**f**, Summary of significant down-regulated (**e**) or up-regulated (**f**) genes for Cas13d- and/or hfCas13d-mediated *PPIA* (g1 or g2), *RPL4*, *CA2* or *PPARG* degradation. **g**, Sites, and relative expression levels of gRNA-dependent off-target transcripts from gRNAs targeting *PPIA* (g1), *PPIA* (g2), *RPL4*, *CA2* or *PPARG* were identified in Cas13d and hfCas13d groups. MM #, mismatch number of off-target sites. Bar value, mean of the biological replicates. **h**, Statistical data analysis from (**g**), among which off-target sites with one or more mismatches were analyzed, each point represents the value of corresponding sample. **i**,**j**, Biological process of significant down-regulated genes induced by Cas13d- or hfCas13d-mediated *PPIA* (**i**) or *RPL4* (**j**) degradation. All values are presented as mean ± s.e.m., n = 3, unless otherwise noted. Two-tailed unpaired two-sample *t*-test. *P < 0.05, **P < 0.01, ***P < 0.001, ns, not significant.

Compared with dCas13d control, we also found numerous off-target changes induced by Cas13d when targeting *RPL4*, *CA2* or *PPARG* transcripts (Fig. 4e-g). Additionally, among those significantly down-regulated genes, targeting transcripts with relatively high expression level (*RPL4*, *PPIA*) in HEK293 cells induced more changes in non-targeted genes than those expressed at low levels (*CA2*, *PPARG*) (Fig. 4e,f), in agreement with the results described above (Figs. 2g,h and 3b,c). Further analysis showed that those down-regulated genes induced by Cas13d targeting with *PPIA* and *RPL4* gRNAs were highly distributed in metabolic and biosynthetic processes (Fig. 4i,j). In addition, most of up-regulated genes induced by Cas13d targeting *RPL4* and *PPIA* were enriched in nucleosome assembly and gene expression pathways, which are related to cellular stress regulation after cleavage events, but not enriched in apoptosis pathways (Supplementary Fig. 9). These results were consistent with previous reports that massive host transcript degradation induced by Cas13 results in retarded growth and dormancy of cells^4, 27, 28, 36^.

Compared with Cas13d, hfCas13d showed marked reduction in the number of down-regulated off-target genes when targeting *PPIA* (9289 vs. 8 for gRNA1; 7271 vs. 19 for gRNA2), *RPL4* (6750 vs. 39), *CA2* (3519 vs. 18), and *PPARG* (1601 vs. 52) (Fig. 4a-f). In addition, hfCas13d also degraded some of the predicted gRNA-dependent off-target genes, although the knockdown efficiency was slightly lower than that of Cas13d (Fig. 4a-d,g,h), indicating mutations in hfCas13d mainly reduced the collateral off-target degradation rather than gRNA-dependent off-target degradation.

Taken together, these RNA-seq results fully confirmed that hfCas13d exhibits high specificity of on-target RNA degradation but no collateral effect.

### No collateral effect of hfCas13d on cell growth, transgenic mice, and *in vivo* somatic editing

To further examine the impact on cellular functions due to collateral effects of Cas13d-mediated RNA degradation *in vivo*, we established doxycycline (dox)-inducible stable cell lines for targeting *RPL4* by Cas13d, hfCas13d, or dCas13d (Fig. 5a). Upon dox treatment, we found that the cell clone carrying Cas13d exhibited significant growth retardation and a notable reduction of *RPL4* transcripts. By contrast, the cell clone carrying hfCas13d exhibited no change in cell growth, and similar reduction of *RPL4* transcripts (Fig. 5b-f). These findings show that collateral effects induced by Cas13d-mediated RNA degradation in HEK293T cells lead to severe growth retardation, and high-fidelity hfCas13d could target specific RNA without affecting cell growth.

**Fig. 5.**
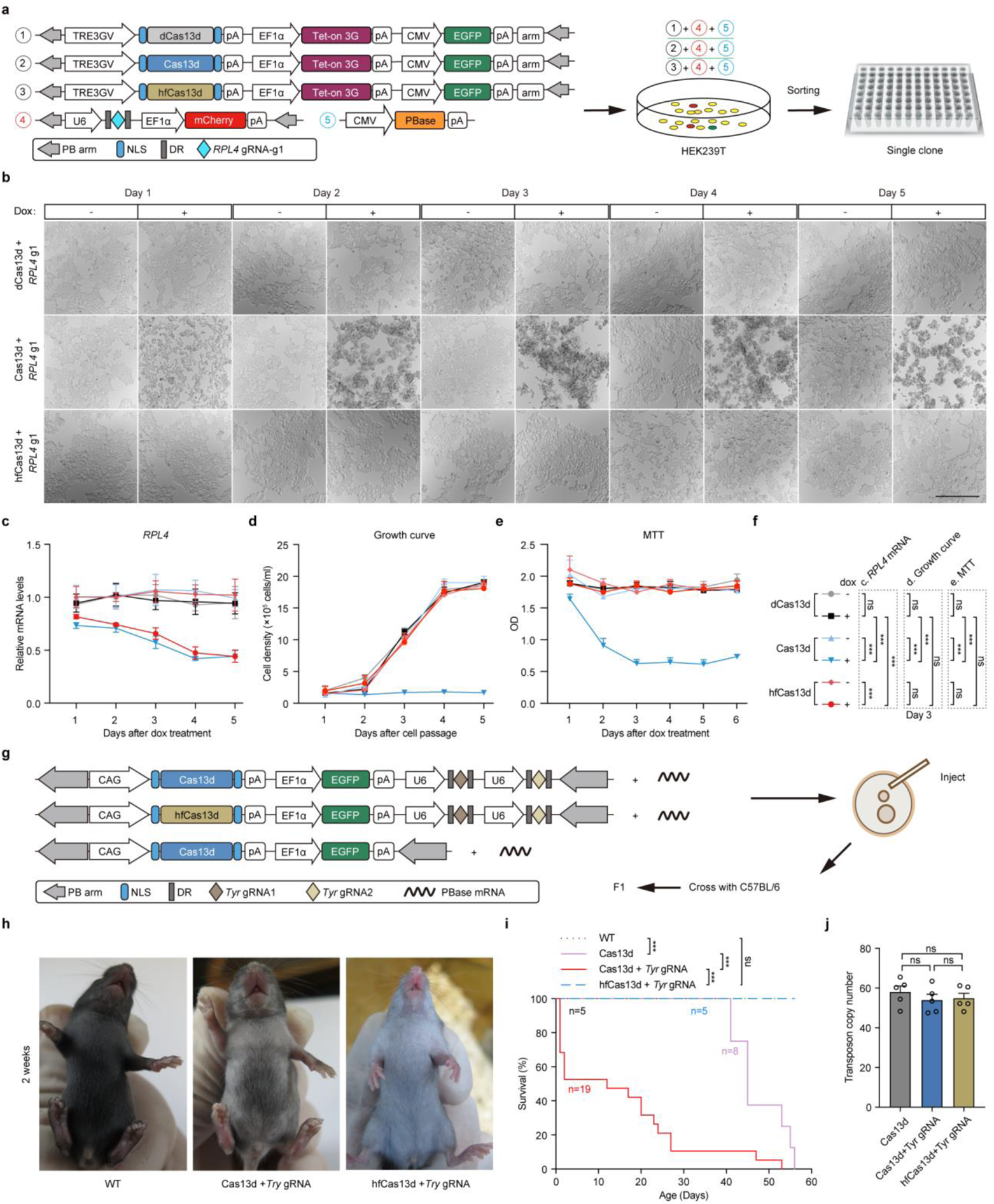
Cellular and *in vivo* consequences of collateral cleavage and their elimination. **a**, Schematic diagram of the dox-inducible dCas13d, Cas13d or hfCas13d and targeting *RPL4* gRNA1 expression system in HEK293T cells. **b**, Representative bright-field images of different cell clones during 5 days after dox treatment. **c-e**, Relative *RPL4* mRNA expression (**c**, n=3), Growth curve (**d**, n=3), and MTT assay (**e**, n=5) of different cell clones with/without dox treatment during 5 or 6 days. **f**, Statistic analysis from (**c-e**). Two-tailed paired two-sample *t*-test. **g**, Schematic diagram of Cas13d or hfCas13d-mediated *Tyr* transcript degradation in mice using the *piggyBac* system. **h**, Representative F1 generation *albino* mice resulting from Cas13d or hfCas13d-mediated *Tyr* knockdown compared to WT (wild-type) mice at 2 weeks. **i**, Kaplan-Meier curve of F1 generation of WT (n = 5) and mice expressing only Cas13d (n = 8), Cas13d with *Tyr* gRNAs (n = 19) or hfCas13d with *Tyr* gRNAs (n = 5). Statistical survival differences were evaluated by two-sided log-rank test. **j**, Copy number of Cas13d or hfCas13d transgenic mice from (**i**), n = 5. All values are presented as mean ± s.e.m.. *P < 0.05, **P < 0.01, ***P < 0.001, ns, not significant.

Moreover, we investigated the collateral effects of Cas13d *in vivo* by generating transgenic mice and examined the RNA-targeting capabilities of Cas13d for endogenous genes, designing gRNAs targeting to *Tyr* (for pigmentation). After co-injection of *PBase* mRNA and *piggyBac* vector containing the gRNAs, Cas13d or hfCas13d, and EGFP reporter into zygotes, we obtained F0 mice carrying Cas13d or hfCas13d and crossed those mice with wild-type C57 mice (Fig. 5g). Two-weeks after birth, white hair phenotype of F1 mice with *Tyr* gRNAs was observed in Bright field, compared to wild-type F1 mice with black hair (Fig. 5h). These findings demonstrated the RNA-targeting activity of Cas13d and hfCas13d *in vivo*. However, compared with the wild types (n = 5), neither mice with only Cas13d (n = 8) nor mice with Cas13d and *Tyr* gRNAs (n = 19) survived more than 8 weeks. Cas13d with non-target gRNA showed collateral activity *in vitro* cleavage assay (Fig. 2i), and thus the lethality of Cas13d mice may be caused by the collateral activity of Cas13d. Moreover, Cas13d with *Tyr* gRNA targeting led to more lethality than without gRNA (Fig. 5i). By contrast, all the mice with hfCas13d and *Tyr* gRNAs (n = 5) survived to adult and showed no abnormality (Fig. 5h,i). The lethality by Cas13d may not due to high copy number of transgenes, as there was no significant difference between Cas13d and hfCas13d transgenic mice (Fig. 5j). To avoid unexpected affects in F0 mice, we constructed conditional expressed mice, in which Cas13d expression could be released by Cre recombinase. We crossed mice containing CAG-LoxP-Stop-LoxP-Cas13d with Cre mice containing *Tyr* gRNAs and mCherry reporter (Supplementary Fig. 10a), and found similar results as that of Cas13d transgenic mice when targeting *Tyr* transcripts (Supplementary Fig. 10b,c). Overall, these finding suggests that Cas13d not only induced off-target lethality, but even worse with targeting gRNA-induced collateral effects, and these adverse effects were eliminated for high-fidelity hfCas13d.

We also examined the collateral effects of Cas13d in somatic cells by intravenous injection of AAV (adeno-associated virus)-packaged Cas13d. PCSK9 is secreted by hepatocytes, and has shown great promise as a candidate of drug targets among all regulators of serum cholesterol^37–39^. AAV8, an efficient liver-targeted gene delivery system^40^, was applied for *in vivo Pcsk9* knockdown. To test the collateral effects of Cas13d targeting, Cas13d or hfCas13d along with *Pcsk9* gRNA1 and gRNA2 were delivered into mice liver through AAV8. The levels of *Pcsk9* mRNA in hepatocytes and PCSK9 protein in serum were significantly reduced in both Cas13d- and hfCas13d-treated mice, compared to those AAV8-EGFP-injected mice 4 weeks after AAV infection (Supplementary Fig. 10d,e). Liver injuries, as indicated by elevated levels of serum aspartate aminotransferase (AST) and alanine aminotransferase (ALT), could be detected in AAV-Cas13d targeting *Pcsk9* mice (Supplementary Fig. 10f,g). By contrast, we did not observe obvious liver injuries in AAV-EGFP or AAV-hfCas13d targeting *Pcsk9* mice (Supplementary Fig. 10f,g). Thus, hfCas13d-mediated RNA knockdown *in vivo* carries significant potential for disease modeling and therapies.

### Generation of hfCas13X by mutagenesis

To generalize the strategy, which we have successfully used to markedly diminish the collateral effects of Cas13d, we chose engineering Cas13X, a newly identified miniature Cas13 protein (775 amino acids) thus more applicable for *in vivo* application^20^. Based on the construction of Cas13d mutant variants and predicted structure of Cas13X (predicted by I-TASSER), we have also developed a mutagenesis library of Cas13X, with mutations mainly located in HEPN1 and HEPN2 domains (Fig. 6a,b), like those of Cas13d. After screening of these mutant variants, we found that Cas13X-M17V6 exhibited low- and high-percentage of EGFP^+^ and mCherry^+^ cells, respectively, indicating high on-target activity but low collateral activity (Fig. 6c,d). Further studies using different EGFP gRNAs showed that Cas13X-M17V6, abbreviated hereafter as hfCas13X, exhibited efficient on-target degradation activity with undetectable collateral effect (Supplementary Fig. 11). In addition, *in vitro* cleavage assay revealed that Casl3X exhibited lower collateral cleavage activity than Cas13d, and hfCas13X showed essentially no collateral cleavage activity (Fig. 6e-g and Supplementary Fig. 12). Moreover, when targeting different endogenous transcripts, hfCas13X exhibited robust degradation of targeted transcripts (Fig. 6h) but with minimal collateral effect (Supplementary Fig. 13).

**Fig. 6.**
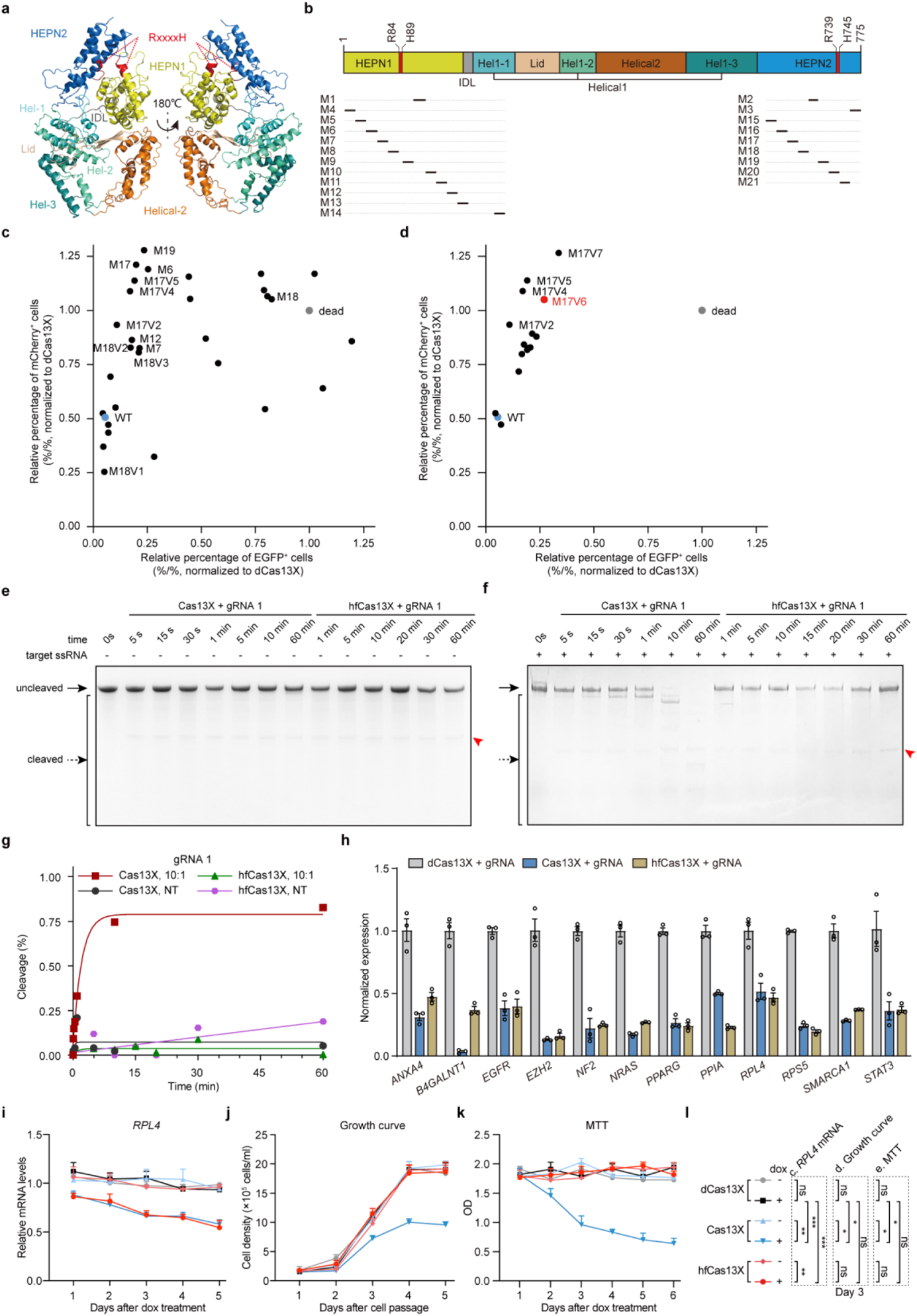
The hfCas13X generation by rational mutagenesis and its efficacy *in vivo*. **a**, Predicted Cas13X structure in ribbon representation. RxxxxHs motifs define the catalytic site, shown as red. **b**, The 21 regions selected for subsequent mutagenesis. **c**,**d**, Quantification of relative percentage of EGFP and/or mCherry positive cells among initially screened Cas13X mutants (**c**) and mutants with different combinations of mutation sites (**d**). Relative percentages of positive cells were normalized to dCas13X (dead Cas13X). WT: wild-type Cas13X. Each dot represents the mean of three biological replicates. **e**,**f**, Representative denaturing gel depicts Cas13X or hfCas13X cleavage activity without target RNA (**e**) or with target RNA (**f**). Red arrows show the band of gRNA. **g**, Quantified time-course data of collateral cleavage by Cas13X or hfCas13X. Exponential fits are shown as solid lines with 10:1 molar ratio of non-target (NT) to target (T) RNA or only with NT RNA. **h**, Relative expression of 12 endogenous transcripts for Cas13X or hfCas13X. dCas13X as vehicle control, n = 3. **g-i**, Relative *RPL4* mRNA expression (**i**), Growth curve (**j**), and MTT assay (**k**) of dCas13X, Cas13X or hfCas13X cell clones with/without dox treatment during 5 or 6 days, n = 3. **l**, Statistic analysis from (**i-k**). Two-tailed paired two-sample *t*-test. All values are presented as mean ± s.e.m.. *P < 0.05, **P < 0.01, ***P < 0.001, ns, not significant.

Similarly, hfCas13X could efficiently degrade endogenous transcripts without affecting cell growth, and eliminate those adverse collateral effects induced by Cas13X (Fig. 6i-l). Cas13X overexpression in transgenic mice did not lead to lethality of mice, but lead to lower body weight than that of wild-type mice and these side effects could be eliminated in hfCas13X transgenic mice (Supplementary Fig. 14). Notably, collateral damage in cell lines and transgenic mice with Cas13d overexpression was more severe than those with Cas13X overexpression, probably due to the high collateral activity of Cas13d.

Together, besides hfCas13d, hfCas13X could also be generated by mutagenesis on HEPN1 or HEPN2 domain of Cas13X, and hfCas13X exhibited minimal collateral effect.

## Discussion

Based on the dual-fluorescence reporter system, transcriptome-wide RNA-seq and cell growth assay, our study presented here has revealed severe Cas13-induced collateral RNA cleavage in mammalian cells, leading to significant cell growth retardation. Therefore, current versions of CRISPR/Cas13 system have a critical deficiency for *in vivo* applications. After a comprehensive mutagenesis screening of Cas13 variants, we obtained several high-fidelity Cas13 variants that retain high on-target RNA cleavage activity but minimal collateral cleavage activity.

Interestingly, we found that many variants exhibited either low dual cleavage activity (upper right in Fig. 2d) or high on-target cleavage activity but low collateral cleavage activity (upper left in Fig. 2d). However, we found no variant exhibiting low on-target cleavage but high collateral cleavage activity (bottom right in Fig. 2d). Previous structural studies have given the target-activated RNA degradation mechanism^30–34^, but the precise dual cleavage mechanism remains unclear.

Our findings suggest a distinct binding mechanism for dual cleavage of Cas13. As depicted in the model for targeted and collateral cleavage (Supplementary Fig. 15), we propose that Cas13 contains two types of separated binding domains proximal to HEPN domains, one specifically for on-target cleavage, and both are required for collateral cleavage. In support of this model, mutations of N1V7, N2V7, N2V8 and N15V4 variants that surround the cleavage site cause steric hindrance or changes in electrostatic interaction (Supplementary Fig. 3), weakening binding between activated Cas13 and promiscuous RNAs without affecting binding between activated Cas13 and target RNAs. Consistent with this model, some variants exhibit elevated dual cleavage activity (bottom left in Fig. 2d and Fig. 6c). These variants with high collateral cleavage activity could facilitate nucleic acid detection^9–11^. Nevertheless, crystal structural model will help to further elucidate the mechanism of high-fidelity Cas13.

Collateral effects have been reported *in vitro* and in bacterial cells, while multiple studies have claimed there is no observable collateral effect in mammalian cells for Cas13a, Cas13b, or Cas13d^5–7^. By contrast, our study and several most recent studies have reported collateral effects of Cas13 proteins in mammalian cells^19, 20, 28, 29^, flies^27^ and mammals. This controversial phenomenon may account for the differences in the expression level of target gene, gRNA sequence, Cas13 orthologs with different collateral activity, the amount of Cas13, and duration of Cas13 expression. In this study, we markedly reduced the collateral activity of Cas13d and Cas13X by mutagenesis, and demonstrated the feasibility of hfCas13d and hfCas13X for efficient on-target RNA degradation with almost no collateral damage in cell lines, transgenic animals, and somatic cells targeting. Although hfCas13d showed mild collateral cleavage activity under conditions that hfCas13d or targeted transcripts expressed at extremely high concentration (Fig. 2g,h and Supplementary Figs. 4 and 7), we could bypass this side effect in most applications.

In short, Cas13 variants with minimal collateral effect we developed are expected to be more competitive for *in vivo* RNA editing and future therapeutic applications.

## Data availability

RNA-seq data are available with the GEO accession number: GSE168246.

## Acknowledgments

We thank Dr. Mu-ming Poo for helpful discussions, insightful comments on this manuscript; Dr. Hui Yang and Beibei Wang from Shanghai Institute of Biochemistry and Cell Biology, Chinese Academy of Sciences for helpful discussions and technical assistance; Optical Imaging facility Y. Wang, Y. Zhang, and Q. Hu, and FACS facility S. Qian, H. Wu and L. Quan of the Center for Excellence in Brain Science and Intelligence Technology, Chinese Academy of Sciences for their technical support. We thank N. Zhong and L. Xie for technical assistance. This work was supported by Chinese National Science and Technology major project R&D Program of China (2017YFC1001302 and 2018YFC2000101), Strategic Priority Research Program of Chinese Academy of Science (XDB32060000), National Natural Science Foundation of China (31871502, 31901047, 31925016, 91957122 and 82021001), Basic Frontier Scientific Research Program of Chinese Academy of Sciences From 0 to 1 original innovation project (ZDBS-LY-SM001), Shanghai Municipal Science and Technology Major Project (2018SHZDZX05), Shanghai City Committee of Science and Technology Project (18411953700, 18JC1410100, 19XD1424400 and 19YF1455100), and International Partnership Program of Chinese Academy of Sciences (153D31KYSB20170059).

## Author contributions

H.T., J.H. and H.Y. jointly conceived the project. H.T., J.H., Q.X., B.H. and X.D. designed and conducted experiments. Y.L. performed bulk RNA-seq analysis. J.H. and X.Y. performed microinjection and counted the mice every day. Q.X., R.Z. and Y.W. performed qPCR assay and participated in FACS. X.D. and D.H. participated in protein purifications and *in vitro* cleavage assays. W.Y performed mouse embryo transfer. Y.L., M.C., Q.W. and M.X. assisted with plasmids construction. Z.W., X.W., C.X., Y.Z., G.L. and K.F. assisted with cell experiments. H.Y. supervised the whole project. H.T., H.Z., and H.Y. wrote the manuscript.

## Methods

### Construction of plasmids

The Cas13d (RfxCas13d) and Cas13a (LwaCas13a) genes and gRNA backbone sequences were synthesized by the HuaGene Company. Then vectors CAG-Cas13d/Cas13a-p2A-GFP, U6-DR-BpiI-BpiI-DR-EF1α-mCherry were generated to knockdown target genes by transient transfection. The gRNA oligos were annealed and ligated into BpiI sites.

### Cell culture, Transfection, and flow cytometry analysis

HEK293T, HEK293 cell lines were purchased from Stem Cell Bank, Chinese Academy of Sciences, and cultured with DMEM (Gibco) supplemented with 10% fetal bovine serum (Gibco), 1% penicillin/streptomycin (Thermo Fisher Scientific) and 0.1 mM non-essential amino acids (Gibco) in an incubator at 37°C with 5% CO_2_.

Cas13 mutants screening was conducted in 48-well plates, and consolidation performed in 24-well plates. The day before transfection, 3×10^4^ HEK293 cells per well were plated in 0.25 mL of complete growth medium. After 12 hours, 0.5 μg plasmids were transfected into cells with 1.25 μg PEI (DNA: PEI = 1:2.5). For 24-well plates, 2×10^5^ cells were plated per well in 0.5 mL of complete growth medium, 0.8 μg plasmids were transfected into HEK293 cells with 2.5 μg PEI. 48 hours after transfection, expression of EGFP and mCherry fluorescence were analyzed by BD FACS Aria III or BD LSRFortessa X-20. Flow cytometry results were analyzed with FlowJo V10.5.3.

### Harvest of total RNA and quantitative PCR

48 hours after transfection, 50,000 of both EGFP and mCherry positive cells were sorted by BD FACS Aria III for RNA extraction. For the groups of mCherry knockdown, total cells of the 12-well plate were collected for RNA extraction. Total RNA was extracted by adding 500 μL Trizol (Invitrogen), 200 μL chloroform to the cells. After centrifuge at 12,000 rpm for 15 min at 4°C, the supernatant was transferred to a 1.5 mL RNase-free tube. 100% isopropanol and 75% alcohol were added to precipitate and purify the RNA. cDNA was prepared using HiScript Q RT SuperMix for qPCR (Vazyme, Biotech) according to manufacturer’s instructions. qPCR reactions were performed with AceQ qPCR SYBR Green Master Mix (Vazyme, Biotech). All the reagents were precooled in advance. qPCR results were analyzed by Roche LC 480 II with −ΔΔCt method.

### Protein purification of Cas13 protein

Cas13 protein purification was performed according to protocol as previously described^41^. The humanized codon-optimized gene for Cas13d, hfCas13d, Cas13X or hfCas13X was synthesized (Huagene) and cloned into a bacterial expression vector (pC013-Twinstrep-SUMO-huLwCas13a from Dr. Feng Zhang laboratory, Plasmid #90097) after the plasmid digestion by BamHI and NotI with NEBuilder HiFi DNA Assembly Cloning Kit (New England Biolabs). The expression constructs were transformed into BL21 (DE3) (TIANGEN) cells. One liter of LB Broth growth media (Tryptone 10.0 g; Yeast Extract 5.0 g; NaCl 10.0 g, Sangon Biotech) was inoculated with ten mL of 12 hours growing culture. Cells were then grown to a cell density A600 of 0.6 at 37°C, and then SUMO-Cas13 proteins expression was induced by supplementing with 0.5 mM IPTG. The induced cells were grown at 16°C for 16-18 hours before harvest by centrifuge (4,000 rpm, 20 min). Collected cells were resuspended in Buffer W (Strep-Tactin Purification Buffer Set, IBA) and lysed using ultrasonic homogenizer (Scientz). Cell debris was removed by centrifugation and the clear lysate was loaded onto StrepTactin Sepharose High Performance Column (StrepTrap HP, GE Healthcare). The non-specific binding protein and contaminants were flowed through. The target proteins were eluted with Elution Buffer (Strep-Tactin Purification Buffer Set, IBA). The N-terminal 6x His/Twinstrep-SUMO tag was removed by SUMO protease (4°C, >20 hours). Then Cas13 proteins were subjected to a final polishing step by gel filtration (Sephacryl 200, GE Healthcare). The purity of >95% was assessed by SDS-PAGE. The proteins were collected and concentrated to about 5mg/ml, and stored at −80°C until use.

### *In vitro* transcription and purification of RNA

The gRNA, target RNA and non-target RNA were transcribed *in vitro*. For gRNA transcription, equimolar concentrations of complementary oligos were mixed in RNase-free water and heated to 95 °C for 5 min, then slowly cooled to 4°C by a touch-down procedure in Biorad PCR instrument. PCR products was used as template for *in vitro* transcription (IVT) using MEGA shortscript T7 kit (Life Technologies) and then purified using MEGA clear kit (Life Technologies). *In vitro* transcription and purification of target /non target RNA were as same as the protocol of gRNA. One difference was that the template for transcription was produced by adding series of primers in a PCR program. All the target RNA, non-target RNA and the gRNAs were eluted in RNase-free water.

### *In vitro* RNA cleavage assay

Cleavage assays were conducted in cleavage buffer (20mM HEPES pH 7.0, 50mM KCl, 2mM MgCl2, 5mM DTT, 5% glycerol) at 37 °C. Cas13 proteins and gRNAs, at a molar ratio of 1:1, were incubated at 37 °C for 20min in cleavage buffer ahead of reactions. In all cleavage assays, reaction mixture without Cas13-gRNA complex was used as non-cleavage blank control. For the concentration course of cleavage assays, 1 μg non-target RNA (NT) were added into the target RNA-Cas13-gRNA mixture (1:1:1) with the final concentrations of Cas13 protein from 17 nM, 34 nM, 68 nM, 136 nM, 550 nM, 1.10 μM, to 2.75 μM. Then the cleavage reaction was processed at 37 °C for 15min. Reactions were terminated by adding 2× RNA loading dye and quenched at 95°C for 10 min. For the time course of cleavage assays with different gRNAs, final concentrations of Cas13 protein and gRNAs were 550 nM.1 μg non-target RNA (NT) was added into the Cas13-gRNA complex, together with target RNA (T) at different molar ratios of NT to T. Then the mixture was incubated for different time periods at 37°C. Samples were analyzed by 15% denaturing TBE-Urea gels. For data analysis, the products were quantified with Image J (National Institutes of Health). Background within each measured substrate was firstly normalized using Image J. The percentage of cleavage was processed as ratio of residual banding intensity relative to un-cleavage band of the blank control. It should be noted that the minimal calculated negative values, if any, due to input variation, were replaced by zero according to the fact that no observed cleavage event occurred. Kinetics data were fitted with a one-phase exponential association curve using Prism (GraphPad).

### RNA-seq and off-targets analysis

For transcriptome sequencing, 35 μg all-in-one plasmids containing Cas13 (dCas13d, Cas13d or hfCas13d), EGFP, mCherry, targeting gRNA for each endogenous gene were transfected into HEK293 cells cultured in 10-cm dishes. 48 hours after transfection, 600,000 dual-positive EGFP^+^/mCherry^+^ (top 15% fluorescent cells) cells were sorted out to make a pool for sequencing by BD FACS Aria III, MoFlo Astrios EQ or Moflo XDP. Total RNA was extracted with TRIZOL-based method, fragmented and reverse transcribed to cDNAs with HiScript Q RT SuperMix for qPCR (Vazyme, Biotech) according to manufacturer’s instructions. RNA-seq library was qualified using Illumina Novaseq 6000 platform in Novogene Co. Ltd or GENEWIZ, respectively. Differential analysis among cell groups (*PPIA* gRNA1, *PPIA* gRNA2, *RPL4* gRNA3, *CA2* gRNA1, *PPARG* gRNA1, and *VEGFA* gRNA1+gRNA2) was performed using a count-based method limma where voom is employed for normalization^42, 43^. BH-adjusted P values (<0.05) and 2-fold-change were used to screen the Significantly expressed genes. Functional analysis was performed using DAVID to identify the enriched biological terms for the significantly changed genes^44, 45^. The biological processes (BP) from Gene Ontology database with FDR (FDR: false discovery rate) values less than 0.05 were selected as significant enriched biological terms.

Off-targets of the gRNAs were predicted using a sequence-based approach. gRNAs in length of 30 bps were first aligned to the human transcriptome (Grch38 cdna from emsembl release 100) using blastn (megablast). Flexible options were set to capture the potential off-targets which could tolerate up to 7 mismatches in this step, i.e., max_target_seqs = 10000, evalue = 10000, word_size = 5, perc_identity = 0.6. Secondly, the candidates were further filtered with the threshold that at least 10 bp perfect match should exist in each alignment.

### Growth curve

Single cell clones with dCas13d, Cas13d, or hfCas13d and *RPL4* gRNA were plated on a 24-well plate at 2×10^5^ cells/mL with or without dox treatment (1 μg/mL). Cell were collected at 24, 48, 72, 96, and 120 hours. Cell number was counted by an automated cell counter (C10311, Invitrogen). Experiments were performed for three replicates.

### Determination of cell proliferation

Cell proliferation was assessed by using a colorimetric thiazolyl blue (MTT) assay. Briefly, single cell clones with dCas13d, Cas13d, or hfCas13d and RPL4 gRNA were treated with or without dox (1 μg/mL) for 0, 24, 48, 72, 96, or 120 hours. Then each group of cells was collected and further plated on a 24-well plate at 2×10^5^ cells/mL with or without dox treatment (1 μg/mL). After an incubation period of 24 hours at 37°C, the tetrazolium salt MTT (Sigma-Chemie) was added to a final concentration of 2 µg/mL, and incubation was continued for 4 hrs. Cells were washed 3 times and finally lysed with dimethyl sulfoxide. Metabolization of MTT directly correlates with the cell number and was quantitated by measuring the absorbance at 550 nm (reference wavelength, 690 nm) using a microplate reader (type 7500; Cambridge Technology, Watertown, MA). Experiments were performed for five replicates.

### Generation of Cas13 transgenic mice

The zygotes used for injection to generate transgenic mice were obtained from the 8-week-old B6D2F1 (C57BL/6 X DBA2J) female mice. ICR female mice (8 weeks of age) were used as recipients. Donor plasmids (100 ng/μl): PB arm-U6 promotor-DR-Tyr gRNA 1-DR-Tyr gRNA 2-DR-CAG promotor-Cas13 (Cas13d, hfCas13d, Cas13X, or hfCas13X)-p2A-GFP-PB arm and PBase mRNA (80 ng/μl) were injected into mouse zygotes. All mice used in this study were housed in a 12 hours light/dark cycle room. All animal experiments were performed and approved by the Animal Care and Use Committee of the Institute of Neuroscience, Chinese Academy of Sciences, Shanghai, China.

### Relative quantification transposon copy number of transgenic mice

qPCR on DNA from mouse tail was used to determine the transposon copy number of individual mouse lines. En2SA DNA amounts were quantified and normalized to beta-actin DNA levels. Transposon copy numbers were determined by normalizing values obtained from Cas13d, hfCas13d, Cas13X or hfCas13X mice to those from wild-type mice that only possessed endogenous En2SA (2 copies).

### Hydrodynamic tail vein injection, serum biochemistry, and hepatocytes isolation

Male C57BL/6 (SLAC laboratory) mice at the age of 8 weeks were used for hydrodynamic tail vein injection. Mice were infected with 1.0×10^11^ transducing units (TU) of AAV in 150 μL PBS by intravenous injection. Mice were sacrificed at 4 weeks post-injection of AAV. Before whole blood collection, mice were fasted for 4 hours. Then whole blood was collected at the time of euthanasia. Whole blood was stood at room temperature for 1 hour and centrifuged at 2000 g for 20 min. Then transfer the serum into new tubes for further analysis. Serum PCSK9 protein were measured with Mouse Proprotein Convertase9/PCSK9 Quantikine ELISA Kit (R&D Systems), according to the manufacturer’s instructions. Serum parameters of liver functions, including alanine aminotransferase (ALT), aspartate transaminase (AST), were measured by automatic biochemical analyzer at the Adicon Clinical Laboratories.inc (Shanghai, China). Mouse primary hepatocytes were isolated by standard two-step collagenase perfusion and purified by 40% Percoll (Sigma) at low-speed centrifugation (1000 rpm, 10 min). Hepatocytes were resuspended in DMEM plus 10% fetal bovine serum (FBS) for FACS, and specific cell populations were used for RNA extraction.

### Statistical analysis

Statistical tests performed by Graphpad Prism 8 included the two-tailed paired two-sample *t*-test, two-tailed unpaired two-sample *t*-test, or the log-rank Mantel-Cox test. The respective statistical test used for each figure is noted in the corresponding figure legends and significant statistical differences are noted as *P < 0.05, **P < 0.01, ***P < 0.001. All values are reported as mean ± s.e.m..

---

**Supplementary Figure 1.**
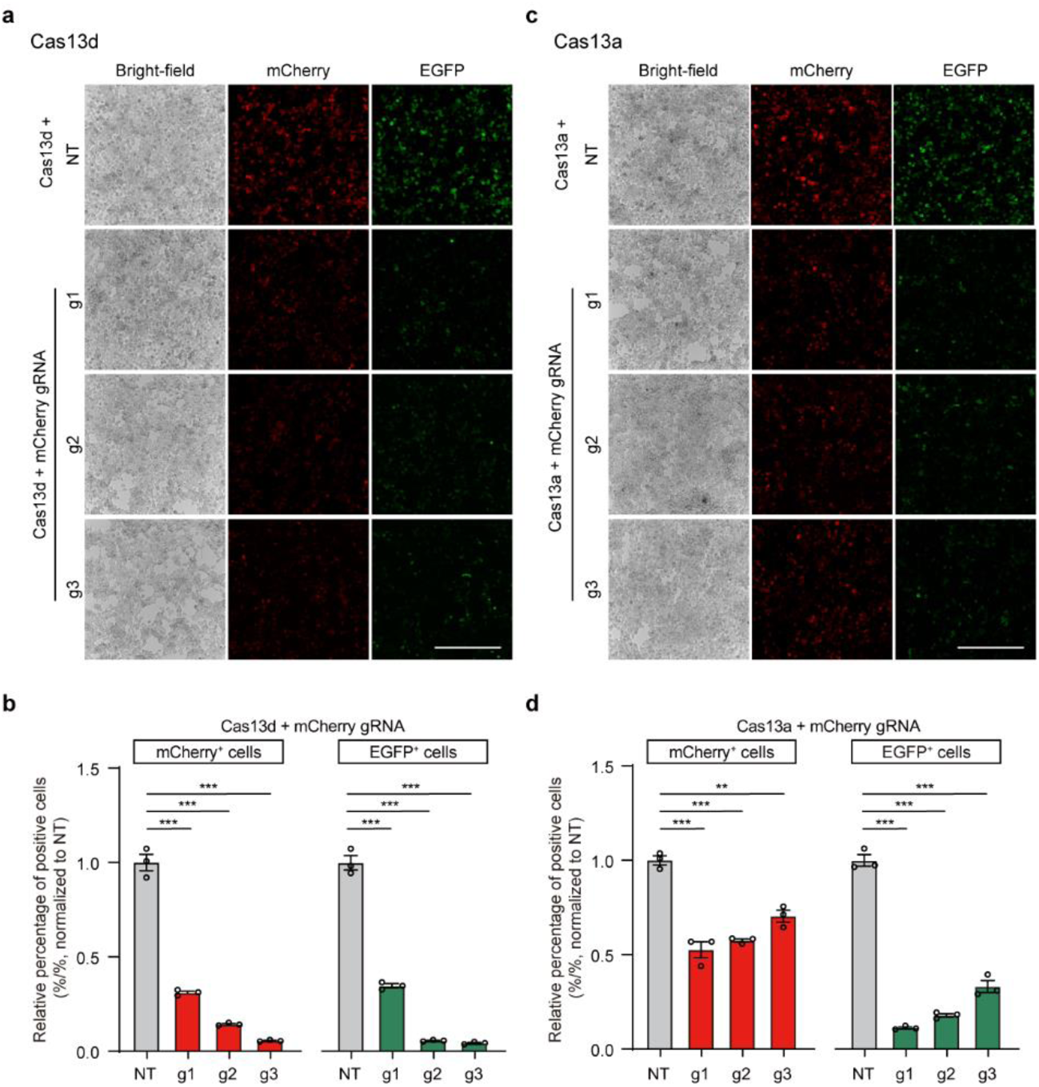
Characteristics of Cas13-mediated mCherry degradation along with collateral effects in HEK293T cells. **a**,**c**, Representative bright-field and fluorescence images of HEK293T cells with mCherry and EGFP RNA degradation by Cas13d (**a**) or Cas13a (**c**) with three different mCherry gRNAs. **b**,**d**, FACS quantitative analysis of relative percentage of EGFP or mCherry positive cells. NT: non-targeting gRNA. All values are presented as mean ± s.e.m., n = 3, unless otherwise noted. Two-tailed unpaired two-sample t-test. *P < 0.05, **P < 0.01, ***P < 0.001, ns, not significant.

**Supplementary Figure 2.**
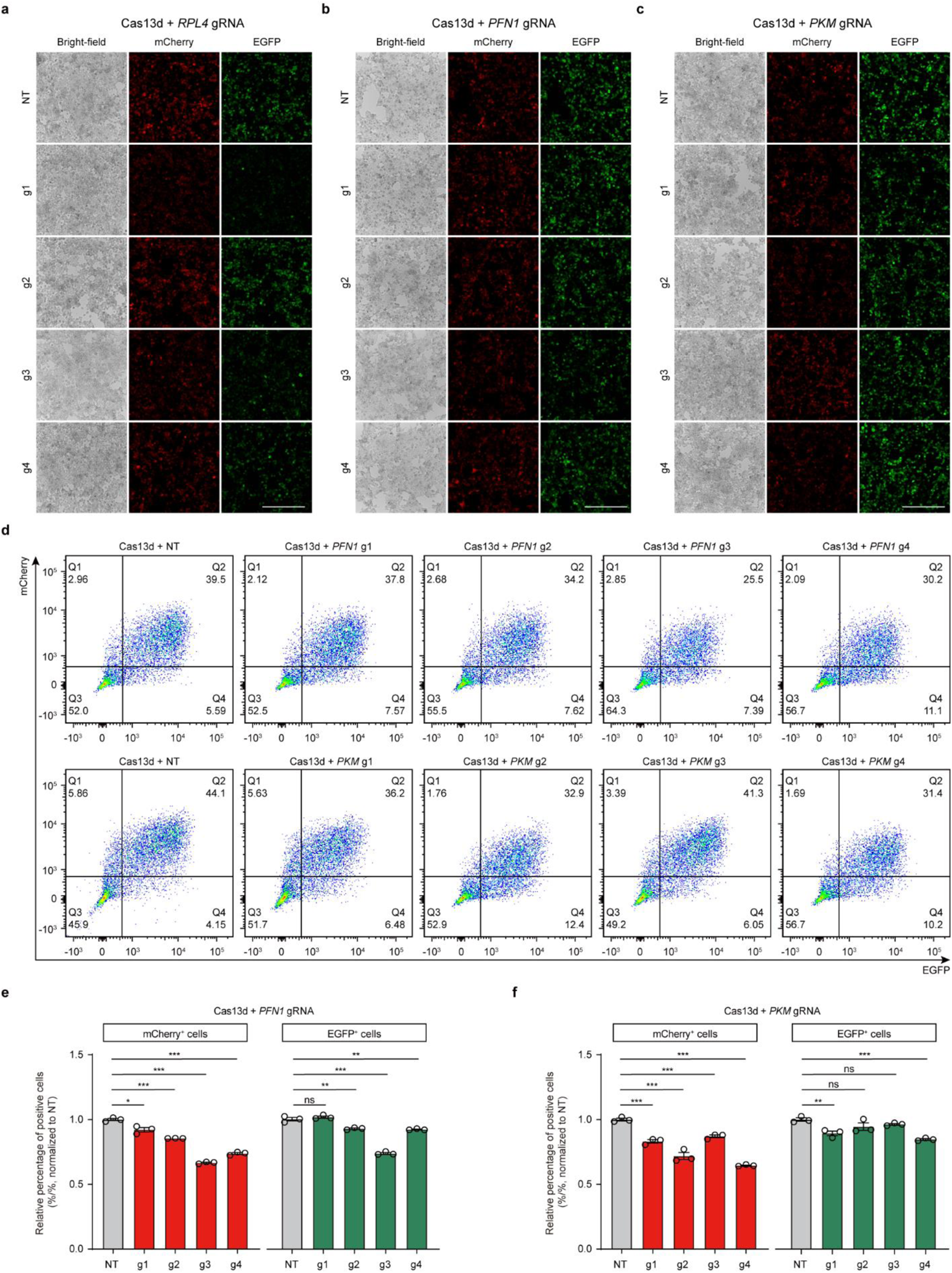
Characteristics collateral effects of Cas13-mediated endogenous transcripts degradation in HEK293T cells. **a**-**c**, Representative bright-field and fluorescence images of cells with reduced mCherry and EGFP fluorescence levels using Cas13d knockdown of three endogenous transcripts (*RPL4*, *PFN1*, or *PKM*), each with four gRNAs. Scale bar, 300 μm. **d**, Representative flow cytometry images of mCherry and EGFP fluorescence intensity from (**b**, **c**). **e**,**f**, Differential decreases of relative percentage of EGFP or mCherry positive cells were induced by Cas13d targeting *PFN1* (**e**), *PKM* (**f**) transcript, with four gRNAs each transcript. NT: non-targeting gRNA. All values are presented as mean ± s.e.m., n = 3, unless otherwise noted. Two-tailed unpaired two-sample t-test. *P < 0.05, **P < 0.01, ***P < 0.001, ns, not significant.

**Supplementary Figure 3.**
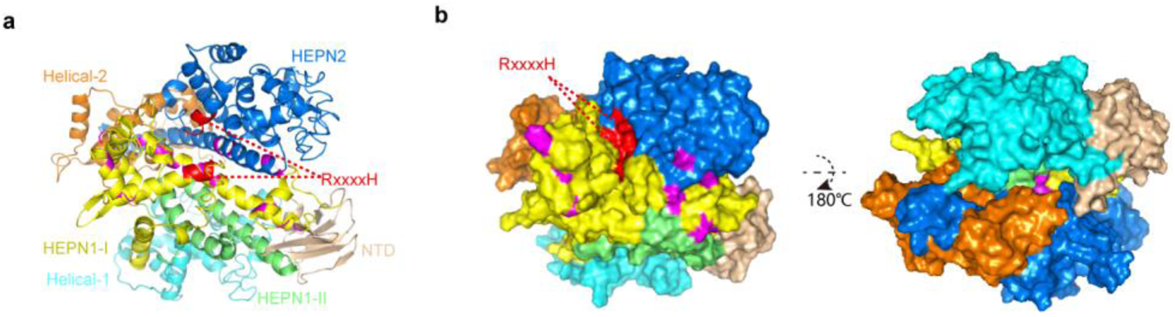
Crystal structure of Cas13d variants. **a**,**b**, Cartoon (**a**) and opposing surface (**b**) view of the predicted overall crystal structure of Cas13d (predicted by I-TASSER). Catalytic sites of the HEPN domains are shown in red, and effective mutant sites are shown in purple.

**Supplementary Figure 4.**
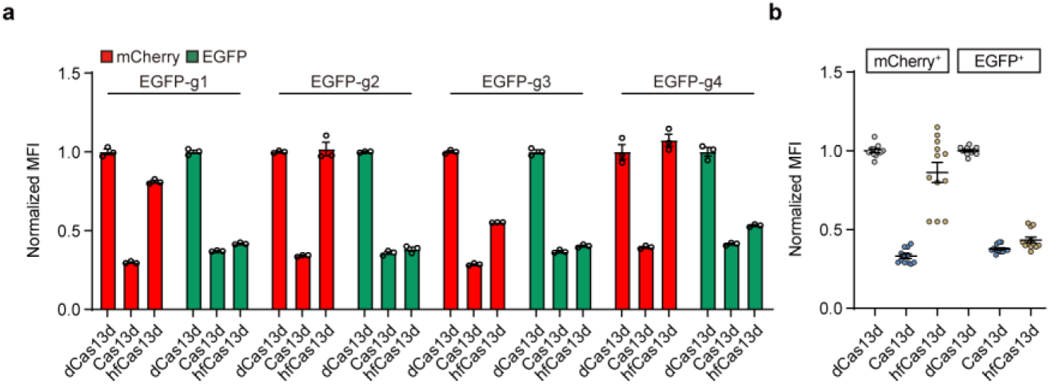
Collateral degradation activity of Cas13d and hfCas13d in HEK293 cells. **a**, Differential changes of normalized MFI were induced by Cas13d, hfCas13d, and dCas13d with EGFP-targeting gRNAs in HEK293 cell lines. **b**, Statistical data analysis for relative degradation efficiency by Cas13d and hfCas13d from (**a**). Normalized MFI, mean fluorescence intensity relative to the dCas13d condition, n = 3. All values are presented as mean ± s.e.m..

**Supplementary Figure 5.**
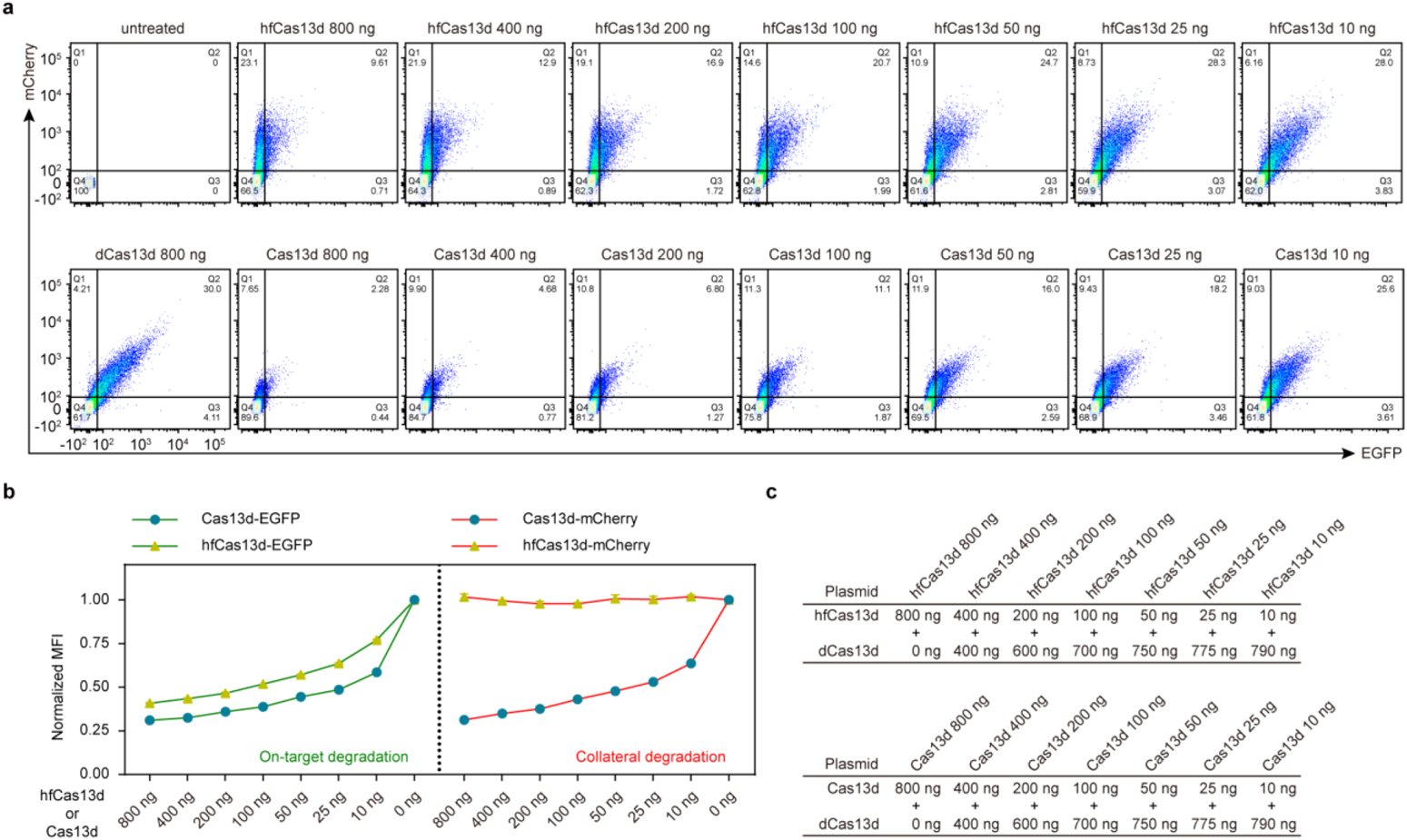
Dose-dependent on-target and collateral degradation activity of Cas13d or hfCas13d. **a**, Representative FACS analysis of mCherry and EGFP degradation induced by transfecting different amount of hfCas13d or Cas13d constructs. The gRNA g2 targeting EGFP was used. **b**, Normalized MFI analysis for (**a**). **c**, A dilution series of Cas13d- or hfCas13d-expressing plasmids mixed with dCas13d-expressing plasmids were transfected. A total of 800 ng plasmids were transfected per well in 24-well plates.

**Supplementary Figure 6.**
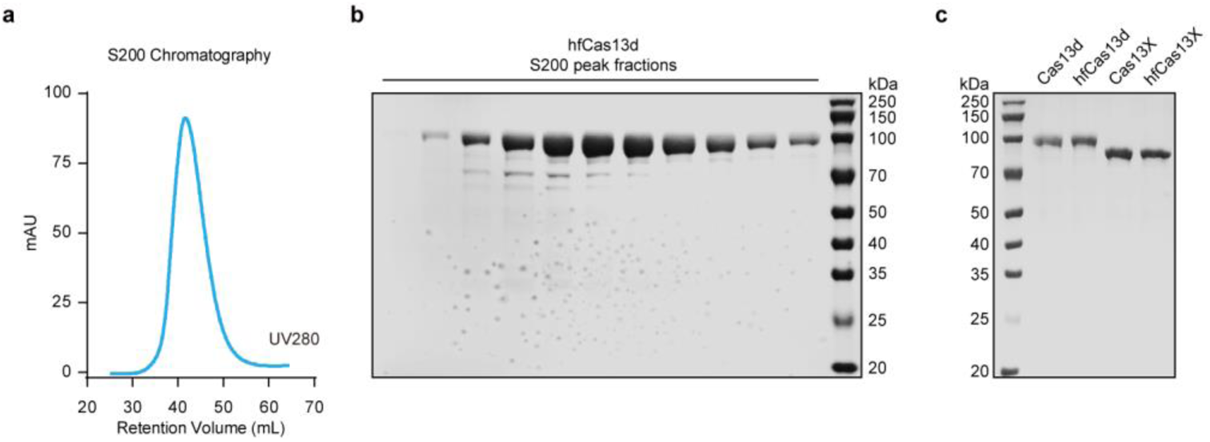
Purification of Recombinant Cas13 Protein. **a**, Chromatogram from Sephacry 200 column for Cas13. The Cas13 fusion protein with N-terminal 6x His/Twinstrep-SUMO tag was purified by successive affinity and size exclusion chromatography. The SUMO-tag was removed by SUMO protease. **b**, SDS-PAGE gel of size exclusion chromatography fractions for Cas13. **c**, SDS-PAGE gel of purified Cas13 proteins.

**Supplementary Figure 7.**
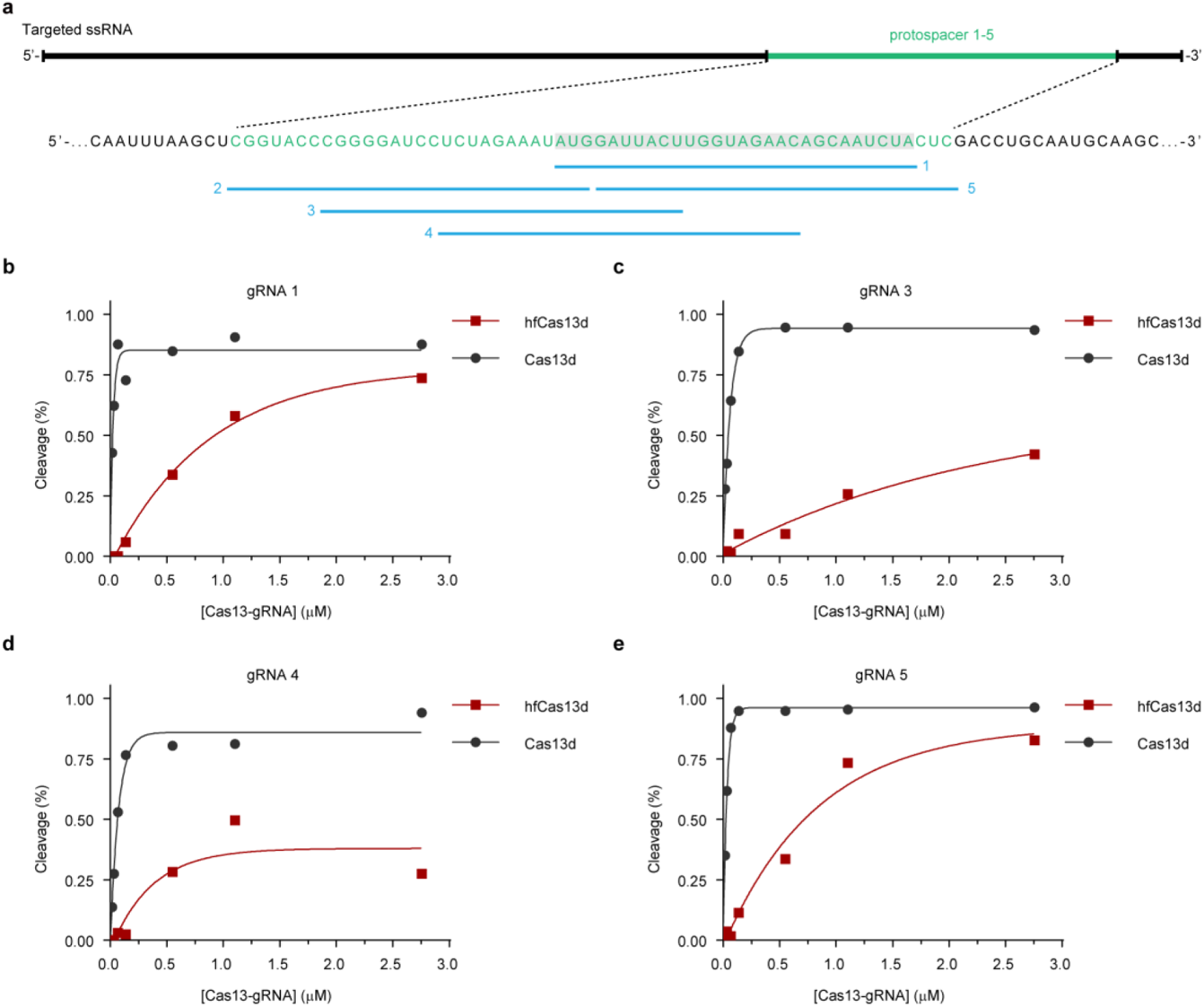
*In Vitro* Characterization of Cas13d and hfCas13d. **a**, Schematic showing the sequence of gRNA spacer and spacer position relative to the complementary ssRNA target. **b-e**, Quantification of collateral cleavage activity of Cas13d or hfCas13d with gRNA 1 (**b**), gRNA 3 (**c**), gRNA 4 (**d**) and gRNA 5 (**e**) in the presence of varying concentrations of Cas13:gRNA complex. Exponential fits are shown as solid lines.

**Supplementary Figure 8.**
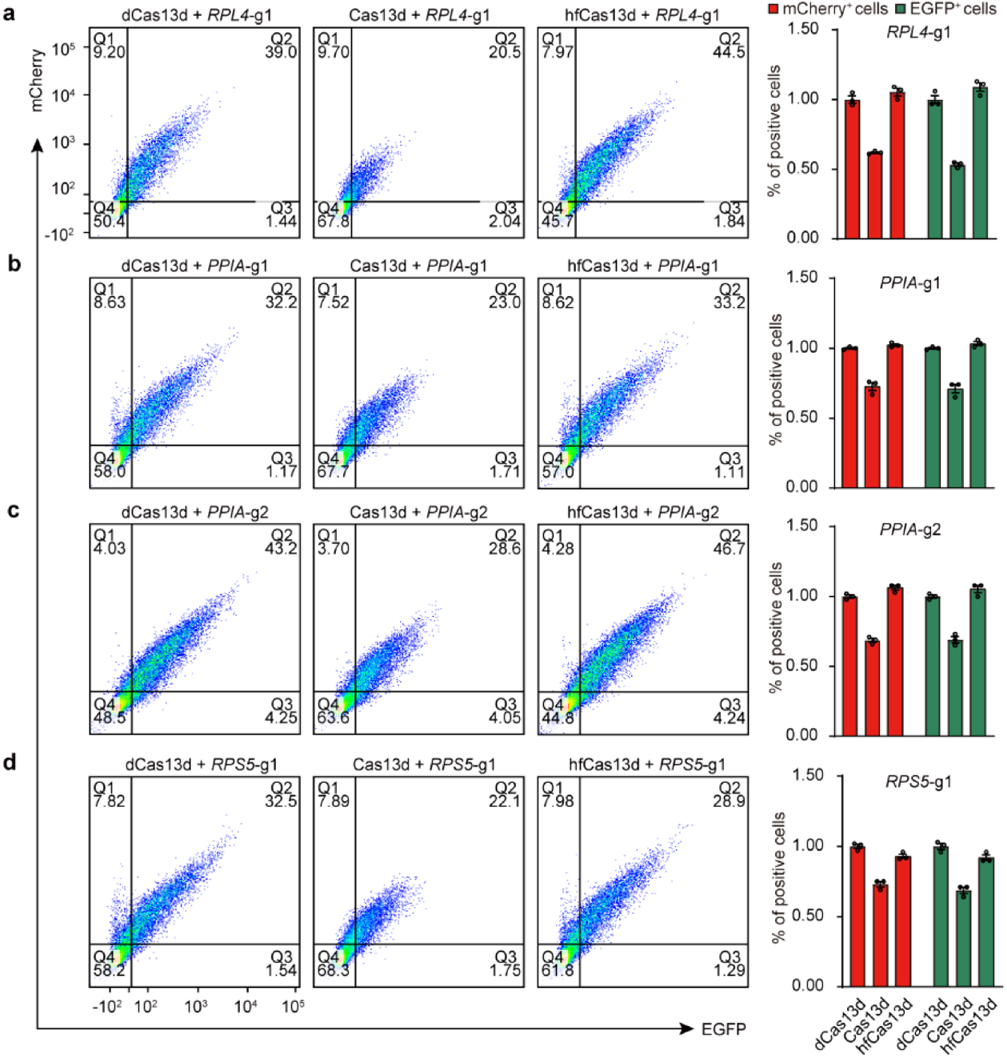
FACS analysis of mCherry and EGFP degradation induced by targeting different endogenous transcripts. Representative FACS analysis of mCherry and EGFP degradation induced by dCas13d, Cas13d, or hfCas13d with gRNA targeting *RPL4* (**a**), *PPIA* (**b**,**c**), and *RPS5* (**d**), respectively, n = 3. CPM, Counts per million. Related to Fig. 3.

**Supplementary Figure 9.**
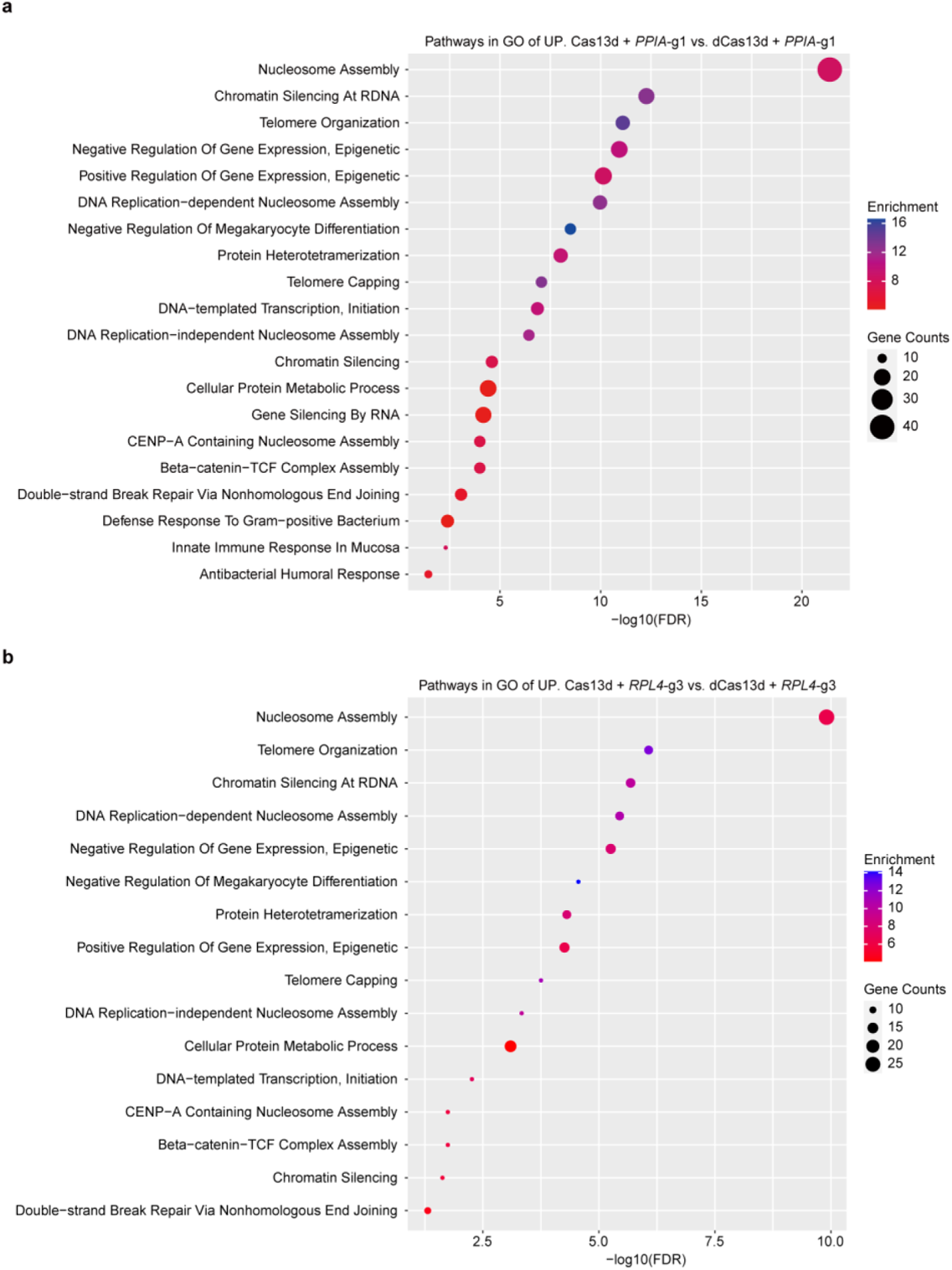
Bulk RNA-seq analysis of genes with differential expression level. Clustering analysis of genes with up-regulation induced by Cas13d targeting *PPIA* (**a**) or *RPL4* (**b**).

**Supplementary Figure 10.**
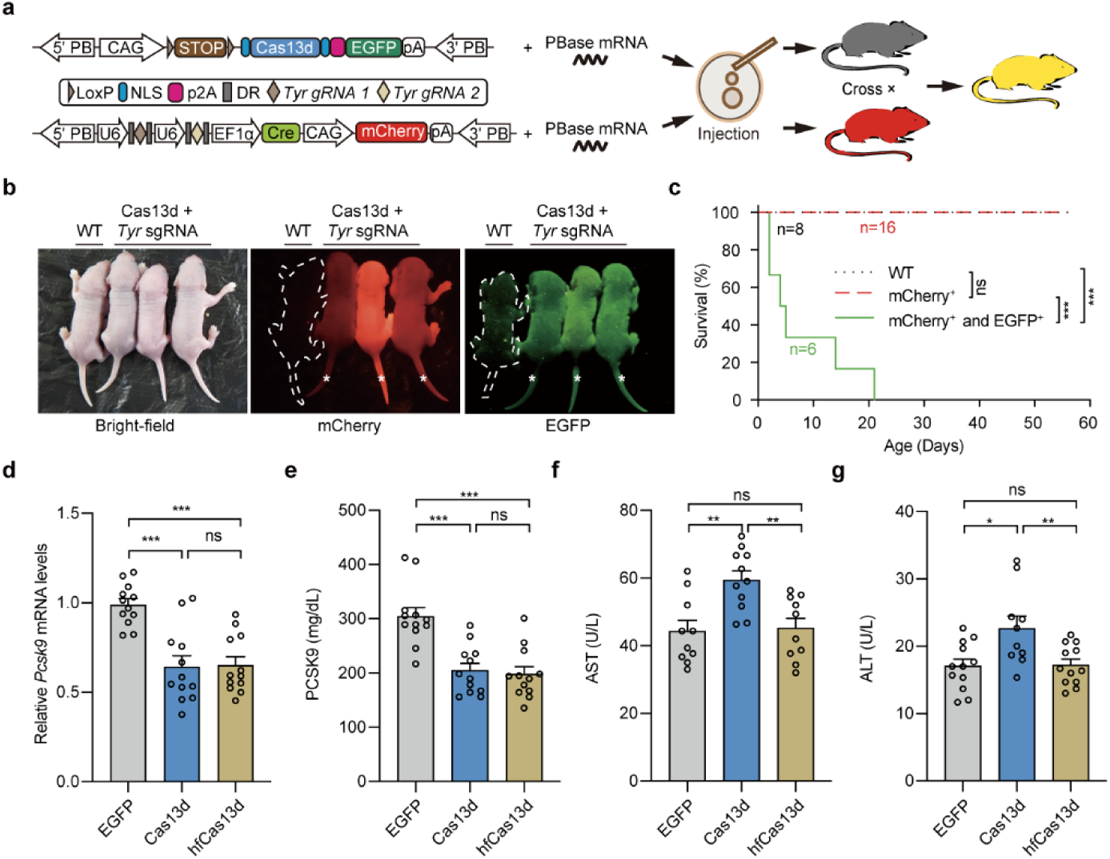
Collateral effects of Cas13d-mediated *Tyr* reduction in mice and *Pcsk9* reduction in the liver. **a**, Schematic diagram of Cas13d-mediated *Tyr* reduction in mice using the *Cre*-LoxP system based on *piggyBac* transposons. **b**, Representative F1 generation mice exhibited EGFP positive along with Cas13d overexpression and mCherry positive along with *Tyr* gRNAs and Cre transposase. White * represents significant expression of EGFP and mCherry. **c**, Survival curve of F1 generation of WT mice, mCherry^+^ transgenic mice and EGFP^+^/mCherry^+^ transgenic mice. Statistical survival differences were evaluated by two-sided log-rank test. **d**,**e**, The levels of *Pcsk9* mRNA in hepatocytes (**d**) and PCSK9 protein in serum (**e**) were quantified at 4 weeks post-injection of AAV, n = 12. **f**,**g**, The levels of serum ALT (**f**) and AST (**g**) were quantified in EGFP-, Cas13d-sgPcsk9-, and hfCas13d-sgPcsk9-injected mice, n = 12. All values are presented as the mean ± s.e.m.. Two-tailed unpaired two-sample *t*-test for (**d-g**). *P < 0.05, **P < 0.01, ***P < 0.001, ns, not significant.

**Supplementary Figure 11.**
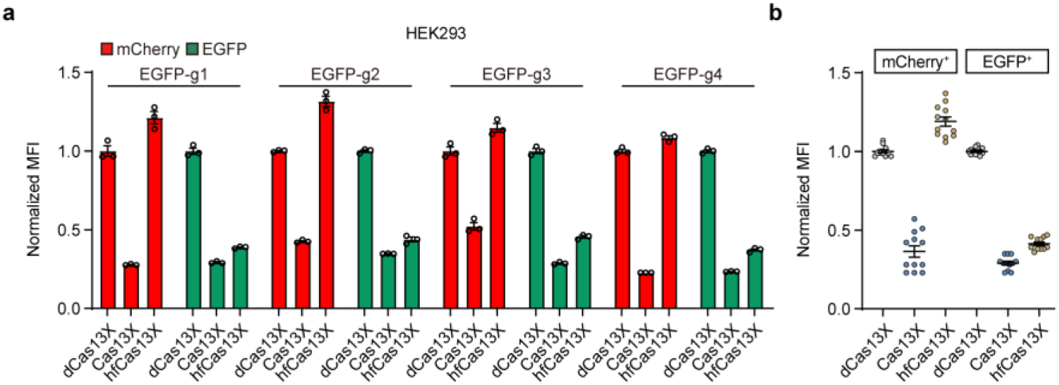
Collateral degradation activity of Cas13X and hfCas13X in HEK293 cells. **a**, Differential changes of normalized MFI were induced by Cas13X, hfCas13X, and dCas13X in HEK293 cell lines. **b**, Statistical data analysis for relative degradation efficiency by Cas13X and hfCas13X from (**a**). Normalized MFI, mean fluorescence intensity relative to the dCas13X condition, n = 3. All values are presented as mean ± s.e.m..

**Supplementary Figure 12.**
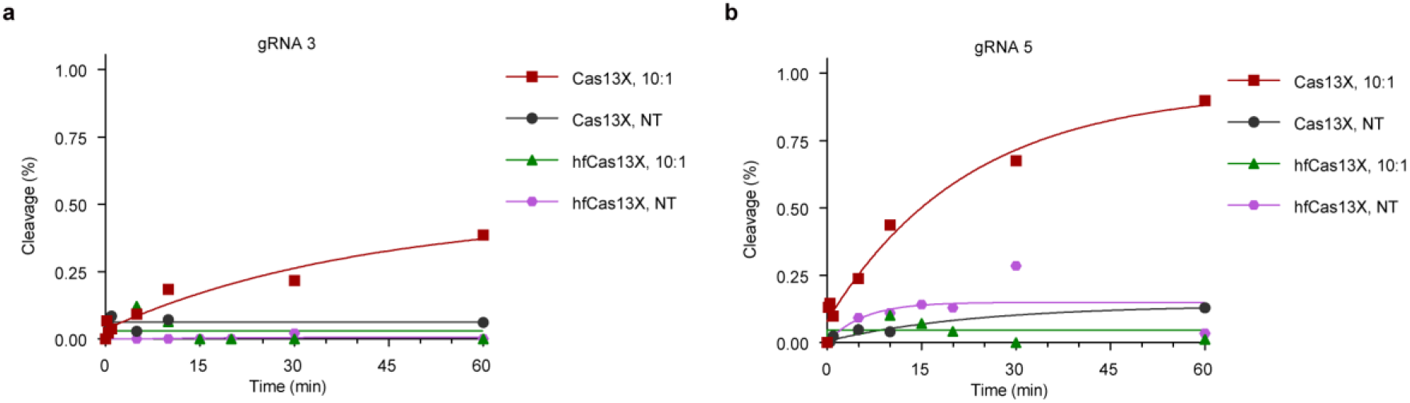
*In Vitro* Characterization of Cas13X and hfCas13X. Quantified time-course data of collateral cleavage by Cas13X or hfCas13X with gRNA 3 (**a**), and gRNA 5 (**b**). Exponential fits are shown as solid lines with 10:1 molar ratio of non-target (NT) to target (T) RNA or only with NT RNA.

**Supplementary Figure 13.**
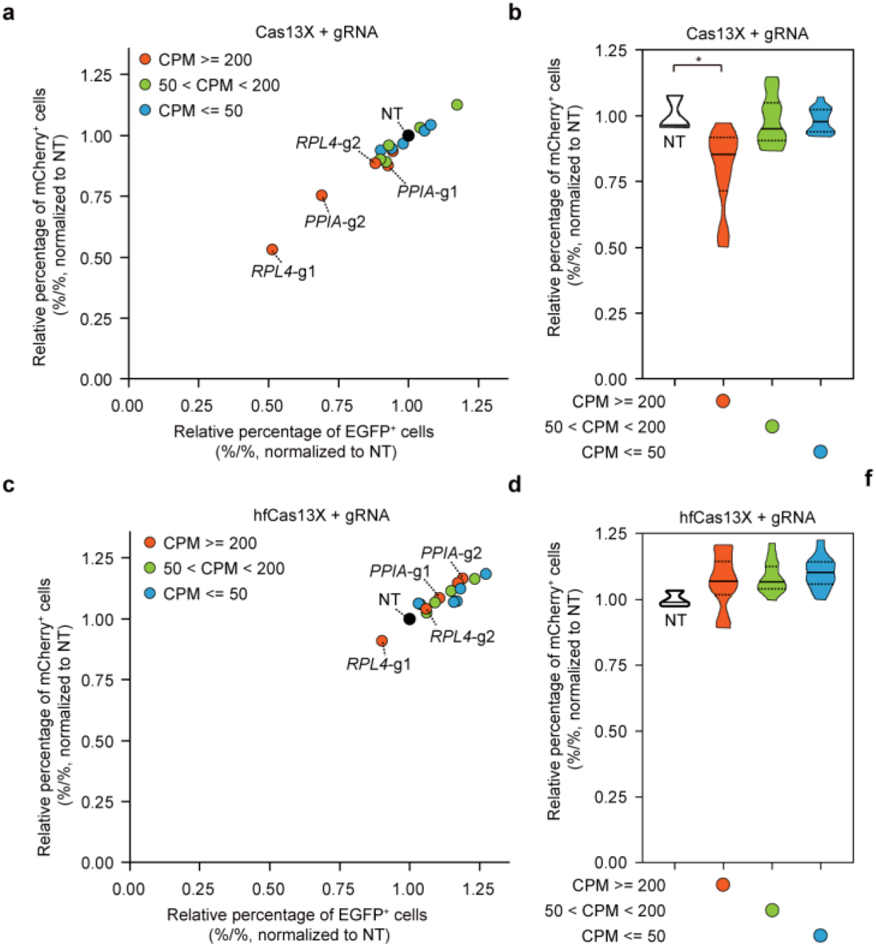
Efficiency and specific interference activity of hfCas13X in HEK293 cells. **a**, Differential reduction of relative percentage of EGFP and/or mCherry positive cells was induced by Cas13X targeting endogenous genes with 15 gRNAs, compared with NT. Each dot represents the mean of three biological replicates for each gRNA targeting endogenous gene with Cas13X. NT: non-targeting gRNA. **b**, Statistical quantification from (**a**). **c**, Differential reduction of relative percentage of EGFP and/or mCherry positive cells was induced by hfCas13X. **d**, Statistical quantification from (**c**). All values are presented as mean ± s.e.m., n = 3, unless otherwise noted. Two-tailed unpaired two-sample *t*-test. *P < 0.05, **P < 0.01, ***P < 0.001, ns, not significant.

**Supplementary Figure 14.**
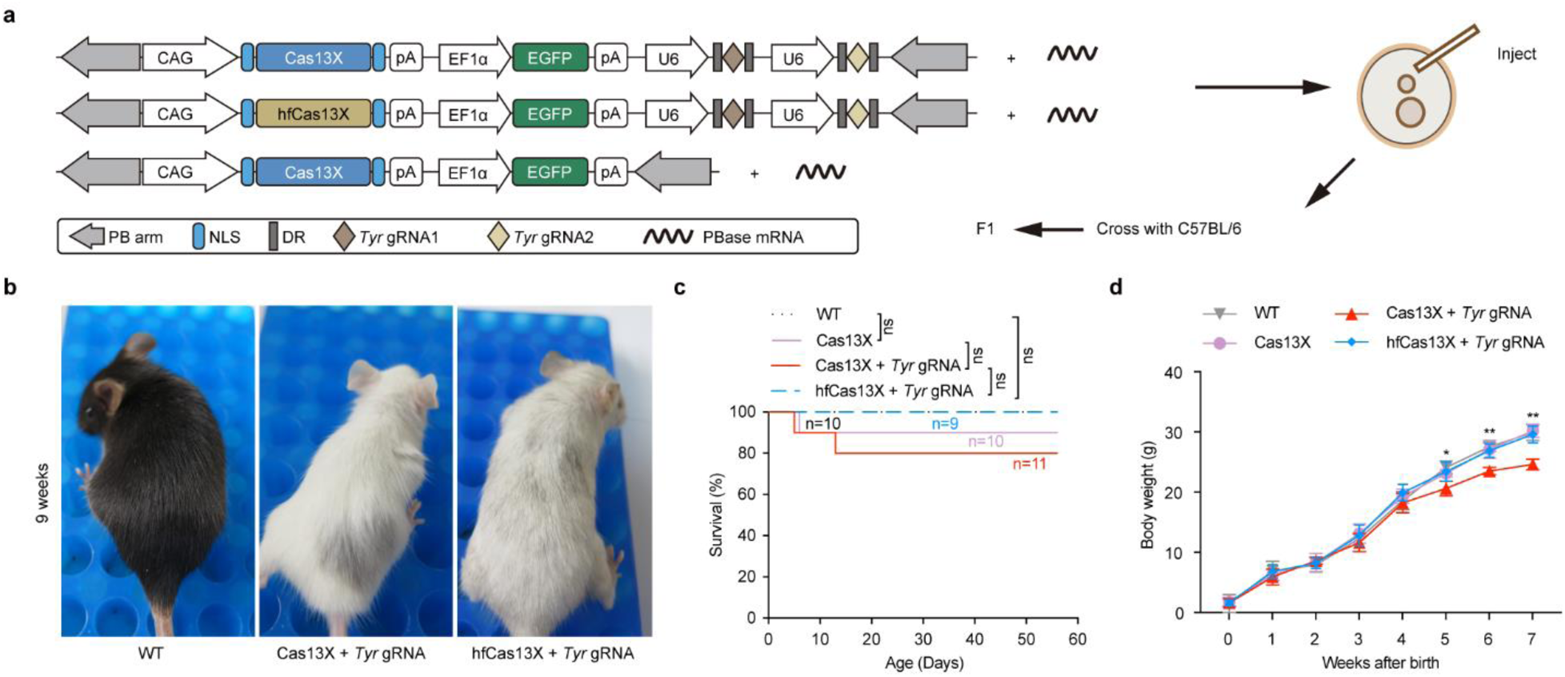
The *in vivo* consequences of collateral cleavage by Cas13X and their elimination. **a**, Schematic diagram of Cas13X or hfCas13X-mediated *Tyr* transcript reduction in mice using the *piggyBac* system. **b**, Representative F1 generation *albino* mouse resulting from Cas13X or hfCas13X-mediated *Tyr* knockdown compared to WT mouse at 9 weeks. **c**, Kaplan-Meier curve of F1 generation of WT mice (n = 10) and mice expressing Cas13X (n = 10), Cas13X with *Tyr* gRNA (n = 11), or hfCas13X with *Tyr* gRNA (n = 9). Statistical survival differences were evaluated by two-sided log-rank test. **d**, Body weight for mice from (**c**). All values are presented as mean ± s.e.m.. Two-tailed unpaired two-sample t-test.*P < 0.05, **P < 0.01, ***P < 0.001, ns, not significant.

**Supplementary Figure 15.**
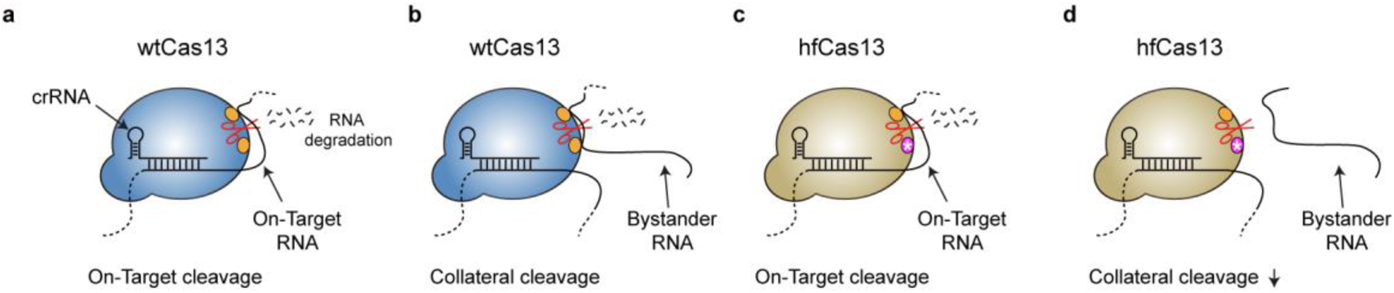
Model of Cas13 on-target and collateral cleavage activity. Once activated by targeted RNA binding, wtCas13 exhibits both on-target cleavage activity and collateral cleavage activity (**a**,**b**), while hfCas13 (e.g., hfCas13d and hfCas13X) with mutant sites maintains undiminished on-target cleavage activity (**c**) but eliminates collateral cleavage activity (**d**).

